# Neuroprotective tissue adaptation induced by IL-12 attenuates CNS inflammation

**DOI:** 10.1101/2022.09.14.506749

**Authors:** Myrto Andreadou, Florian Ingelfinger, Donatella De Feo, Ekaterina Friebel, Selma Tuzlak, Teresa M.L. Cramer, Bettina Schreiner, Pascale Eede, Shirin Schneeberger, Maria Geesdorf, Frederike Ridder, Christina Welsh, Daniel Kirschenbaum, Shiva K. Tyagarajan, Melanie Greter, Frank L. Heppner, Sarah Mundt, Burkhard Becher

## Abstract

IL-12 is a well-established driver of type 1 immune responses. Paradoxically, in several autoimmune conditions including neuroinflammation, IL-12 reduces pathology and exhibits regulatory properties. Yet, the mechanism and the involved cellular players behind this immune regulation remain elusive. To identify the IL-12-responsive elements which prevent immunopathology, we generated mouse models lacking a functional IL-12 receptor either in all cells or in specific populations within the immune or central nervous system (CNS) compartments, and induced experimental autoimmune encephalomyelitis (EAE), which models human Multiple Sclerosis (MS). This revealed that the CNS tissue-protective features of IL-12 are mediated by cells of the neuroectoderm, and not immune cells. Importantly, sections of brain from patients with MS show comparable patterns of expression, indicating parallel mechanisms in humans. By combining spectral flow cytometry, bulk and single-nucleus RNA sequencing, we uncovered an IL-12-induced neuroprotective adaption of the neuroectoderm critically involved in maintaining CNS tissue integrity during inflammation.

## Main

MS is a chronic inflammatory disease of the CNS characterized by cytokine dysregulation, demyelination and neuronal loss^1^. The inflammatory lesions in MS result from type-1 immunity, which drives lymphocyte and monocyte infiltration and phagocyte-mediated immunopathology^2^. The archetypical inducer of type 1 immunity is the cytokine IL-12, produced by antigen-presenting cells^3^.However, with the discovery of the closely related cytokine of the same superfamily, namely IL-23^4^, we and others found that IL-23 plays a non-redundant role in the development of CNS inflammation in preclinical models of MS^5,6^. Surprisingly, these studies also discovered that IL-12 ameliorated the disease^7,8^ and has therefore a regulatory immunosuppressive role in neuroinflammation. IL-12 and IL-23 are heterodimers harbouring the unique subunits p35 and p19 engaging with IL-12Rβ2 and IL-23R respectively, while sharing the common p40 subunit, which binds IL-12Rβ1. Dual targeting of IL-12 and IL-23 signalling, however, has not shown efficacy in a phase two clinical trial in MS therapy^9^, potentially due to interfering with two opposing underlying disease mechanisms simultaneously. In fact, higher *IL12RB2* expression has been correlated with a lower risk of relapse in RRMS patients and better response to MS therapy^10,11^. Thus, IL-12 may attenuate neuroinflammation not only in mice but in human MS, too. The mechanism behind IL-12-mediated tissue protection and immune regulation in MS has never been resolved.

To identify the mechanistic underpinnings of how IL-12 limits immunopathology and neuroinflammation, we generated an *Il12rb2* conditional knockout mouse line. We found that, in addition to broad expression of the IL-12 receptor (IL-12R) complex across leukocytes, IL-12R was also expressed in cells of the CNS neuroectoderm, specifically in neurons and oligodendrocytes, of both mice and humans. We discovered that the conditional ablation of the *Il12rb2* allele within the neuroectoderm alone phenocopied the hypersusceptibility to EAE observed in germline *Il12rb2* knockout mice. Here, we describe an IL-12-driven neuroprotective loop implicating autocrine and paracrine survival and trophic factor release by distinct neuronal cell populations in neuroinflammation.

## Results

### Hematopoietic cells are dispensable for IL-12-mediated tissue protection in EAE

To systematically interrogate the mechanism by which IL-12 limits immunopathology in MS, we first generated a conditional strain of IL-12-specific receptor subunit *b2: Il12rb2*^*f*/12^. We crossed *Il12rb2*^fl/fl^ mice with *CMV^Cre^* mice to generate mice lacking *Il12rb2* in all tissues, hereafter termed *Il12rb2*^del/del^ mice (Fig. 1a). As evidence of successful *Il12rb2* ablation, leukocytes from *Il12rb2*^del/del^ mice were unable to phosphorylate STAT4 in response to exogenous IL-12 *in vitro* (Extended Data Fig. 1a). Moreover, these mice were hypersusceptible to myelin oligodendrocyte glycoprotein (MOG)-induced EAE, as previously reported in mice lacking functional IL-12^7,8^ (Fig. 1b-c). While the time to clinical disease onset was comparable between mice with or without *Il12rb2* (Fig. 1b), the absence of IL-12 signalling was associated with significantly more severe disease development (Fig. 1b, Extended Data Fig. 1b).

**Figure 1:**
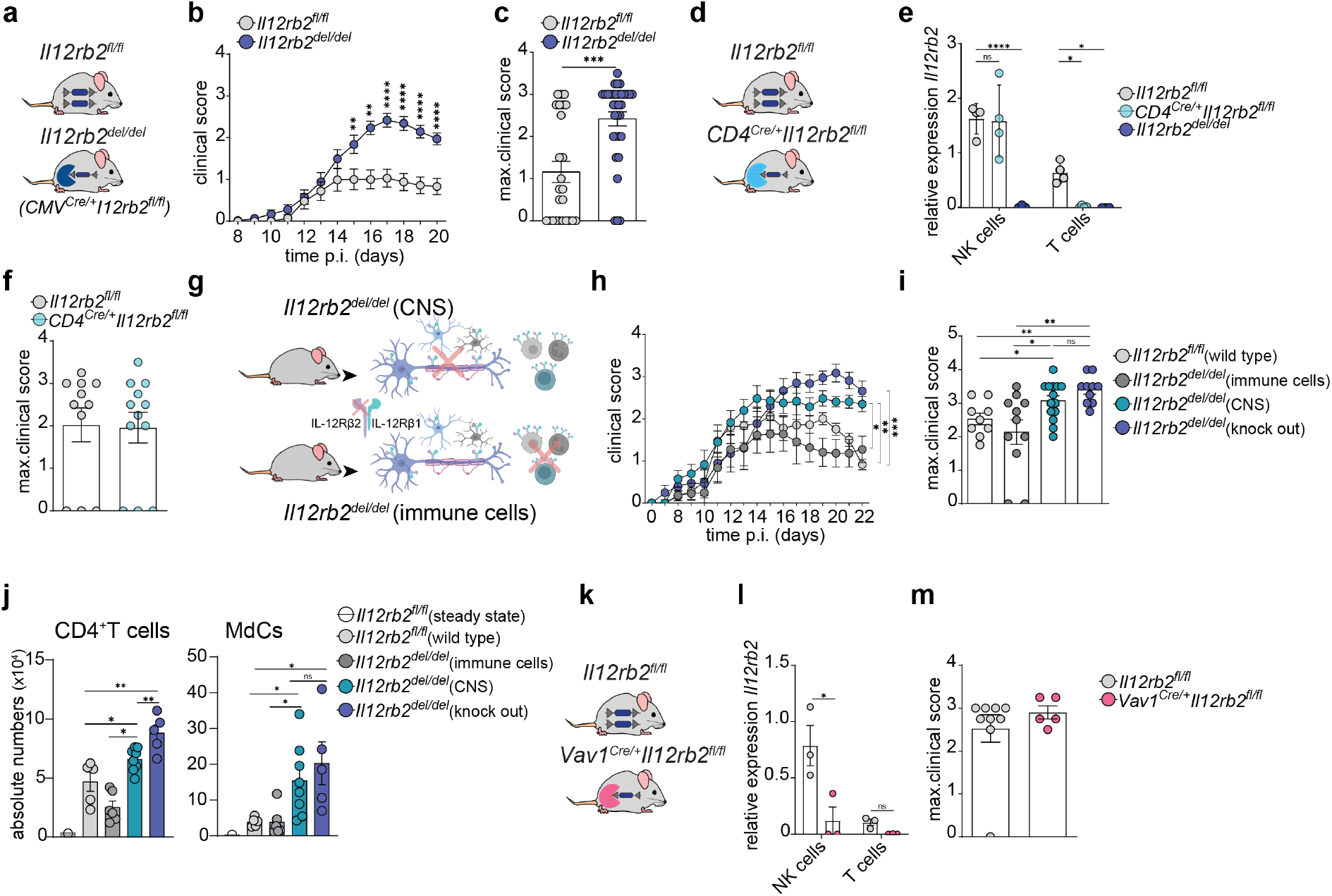
Hematopoietic cells are dispensable for the tissue-protective features of IL-12 in EAE. (a) *CMV^Cre^* mice were crossed to *Il12rb2^fl/fl^* mice to generate *Il12rb2^del/del^* mice, which lack *Il12rb2* in all tissues. (b) Clinical disease course and (c) maximum clinical scores of MOG-CFA induced EAE in *Il12rb2^del/del^* (n=33) and *Il12rb2^fl/fl^* mice (n=25). Data pooled from four independent experiments using both male and female mice. (d) *CD4*^*Cre*/+^ mice were crossed to *Il12rb2^fl/fl^* mice to generate *CD4*^*Cre*/+^*Il12rb2^fl/fl^* mice, which lack *Il12rb2* in αβ T cells. (e) *Il12rb2* mRNA relative expression level in NK cells and CD3^+^ T cells isolated by FACS from the brains of *CD4*^*Cr*e/+^*Il12rb2^fl/fl^* (n=4), littermate controls (n=4) and *Il12rb2^del/del^* (n=3) mice at 18 days post immunization (dpi). Data represent one experiment. (f) Individual maximal EAE scores of MOG-CFA induced EAE in *CD4*^*Cre*/+^*Il12rb2^fl/fl^* (n=12) and littermate controls (n=11). Data are pooled from two independent experiments. (g) Illustration depicting IL-12R expression in bone marrow chimeras (created with Biorender.com). (h) Clinical disease course of MOG-CFA induced EAE in *Il12rb2^del/del^* and *Il12rb2^fl/fl^* mice, lethally irradiated at 6 weeks of age and reconstituted with bone marrow isolated either from *Il12rb2^del/del^* or *Il12rb2^fl/fl^* mice of the same age. Data pooled from two independent experiments with n=9-14 per group. (i) Maximum EAE scores of mice in (h). (j) Absolute numbers of CD4^+^ T cells (CD45^+^CD44^+^CX3CR1^-^LY6G^-^Ly-6C^-^TCR-beta^+^CD8^-^CD4^+^) and MdCs (CD45^+^CD44^+^CX3CR1^-^LY6G^-^TCR-beta^-^NK1.1^-^SiglecF^-^CD11b^+^Ly-6C^+^MHCII^+^) in the CNS at 23 dpi as measured by spectral flow cytometry. The data are representative of two independent experiments with n=4-5 per group. (k) *Vav1*^*Cre*/+^ mice were crossed to *Il12rb2^fl/fl^* mice to generate *Vav1^Cre/+^Il12rb2^fl/fl^* mice, which lack *Il12rb2* in hematopoietic cells. (l) *Il12rb2* mRNA expression of NK cells and CD3^+^ T cells isolated by FACS from the whole brains of *Vav1^Cre/+^Il12rb2^fl/fl^* (n=3) mice and littermate controls (n=3) at 19 dpi. (m) Maximal EAE scores in *Vav1*^*Cre*/+^*Il12rb2^fl/fl^* (n=5) and littermate controls (n=9). The data are representative of two independent experiments. Each symbol represents one mouse. Data are shown as mean ± SEM. In (b), (e), (h), (l) statistical significance was evaluated by two-way ANOVA with Bonferroni’s post hoc test; in (c), (f), (i) and (m) by a non-parametric Mann-Whitney test; an unpaired Student’s *t* test was used to compare the means in (j).* P < 0.05, ** P < 0.01,*** P < 0.001, ****P < 0.0001, ns=not significant.

As helper T cells (T_H_) are the main drivers of MOG-induced EAE^13^, we asked whether deleting the IL-12R across aβ T cells could recapitulate the EAE hypersusceptibility seen in *Il12rb2^del/del^* mice. We crossed *Il12rb2^fl/fl^* mice with *CD4*^*Cre*/+^ mice (Fig. 1d), generating progeny in which *Il12rb2* was deleted in T cells (Fig. 1e, Extended Data Fig. 1c). By analyzing IL-12-induced phosphorylation of STAT4 and qPCR (Fig. 1f, Extended Data Fig. 1c), we confirmed the deletion of *Il12rb2* in T cells. Surprisingly, both *CD4^Cre/+^Il12rb2^fl/fl^* mice and *CD4*^+/+^*Il12rb2^fl/fl^* littermates developed EAE of similar clinical severity (Fig. 1f, Extended Data Fig. 1d) suggesting that T cells are not responsible for the IL-12-mediated tissue protection. Similarly, using the *Nkp46*^*Cre*/+^*Il12rb2*^fl/fl^ we excluded a role for *Il12rb2-bearing* NK cells in the immunoregulatory circuit (Extended Data Fig. 1e-h) thereby eliminating the most obvious targets within the lymphocyte compartment.

To understand whether IL-12 acts as a regulator of autoimmune CNS inflammation in another type of immune cell or directly on CNS cells, we generated bone marrow chimeras in which *Il12rb2* was expressed either in the adult hematopoietic stem cell (aHSC)-derived systemic immune compartment or in the CNS (Fig. 1g). When we induced EAE in these mice, we found that the loss of IL-12-sensing in the radioresistant CNS neuroectoderm (i.e. neurons, oligodendrocytes and astrocytes) and microglia phenocopied the *Il12rb2^del/del^* strain (Fig. 1h-i). Mice lacking *Il12rb2* expression in CNS-intrinsic cells also showed significantly more infiltration of CD4^+^T cells and monocyte-derived cells (MdCs) into their CNS (Fig. 1j). Further, depleting *Il12rb2* from all hematopoietic cells by crossing the *Vav1^Cre^* with the *Il12rb2^fl/fl^* mouse strain, allowed to eliminate all hematopoietic cells (including CNS-resident microglia^14^) from the list of potential candidates for the IL-12-mediated protective phenotype in EAE (Figure 1k-m, Extended Data Fig. 1i-j). Taken together, these data illustrate that the protective features of IL-12 were mediated by CNS intrinsic cells independent of immune cells.

### Neurons and oligodendrocytes are molecularly equipped to sense IL-12 in mouse and human

We next sought to capture and localize potential sensors of IL-12, resident to the murine CNS. Leveraging the capacity of the *Il12rb2* knockout-first allele to act as a reporter^12^, we used immunostaining for β-gal to detect cells expressing IL-12Rβ2 protein within the steady-state CNS, coupled with multiplexed RNA fluorescence *in situ* hybridization (RNAscope^®^) for *Il12rb1* and *Il12rb2* mRNA to capture both components of the IL-12R. IL-12R was expressed in the neuroectoderm, specifically in neurons and oligodendrocytes, already in the healthy brain (Fig. 2a-h, Extended Data Fig. 2a-b), which is consistent with recently published single-nucleus RNA sequencing (snRNA-seq) atlases of the steady-state, adult mouse cerebellum^15^ and hippocampus^16,17^. In fact, both β-gal/IL-12Rβ2 immunoreactivity (Fig. 2a-h, Extended Data Fig. 2a-b) and *Il12rb2* mRNA transcripts (Fig. 2i-j) were localized to NeuN^+^- and *Rbfox3/Map2-positive* neuronal cells, respectively. Of note, calbindin^+^ Purkinje cells in the cerebellum showed particularly strong IL-12Rβ2 expression (Fig. 2b). The β-gal-positive signal was also present in CC1^+^ myelin-forming and mature oligodendrocytes (Fig. 2g). Notably, *Il12rb1* mRNA was more strongly expressed in *Sox-10*-positive oligodendrocytes (Fig. 2i-j). Neither astrocytes nor microglia expressed the IL-12R subunits, suggesting their inability to sense IL-12 (Fig. 2f, j).

**Figure 2:**
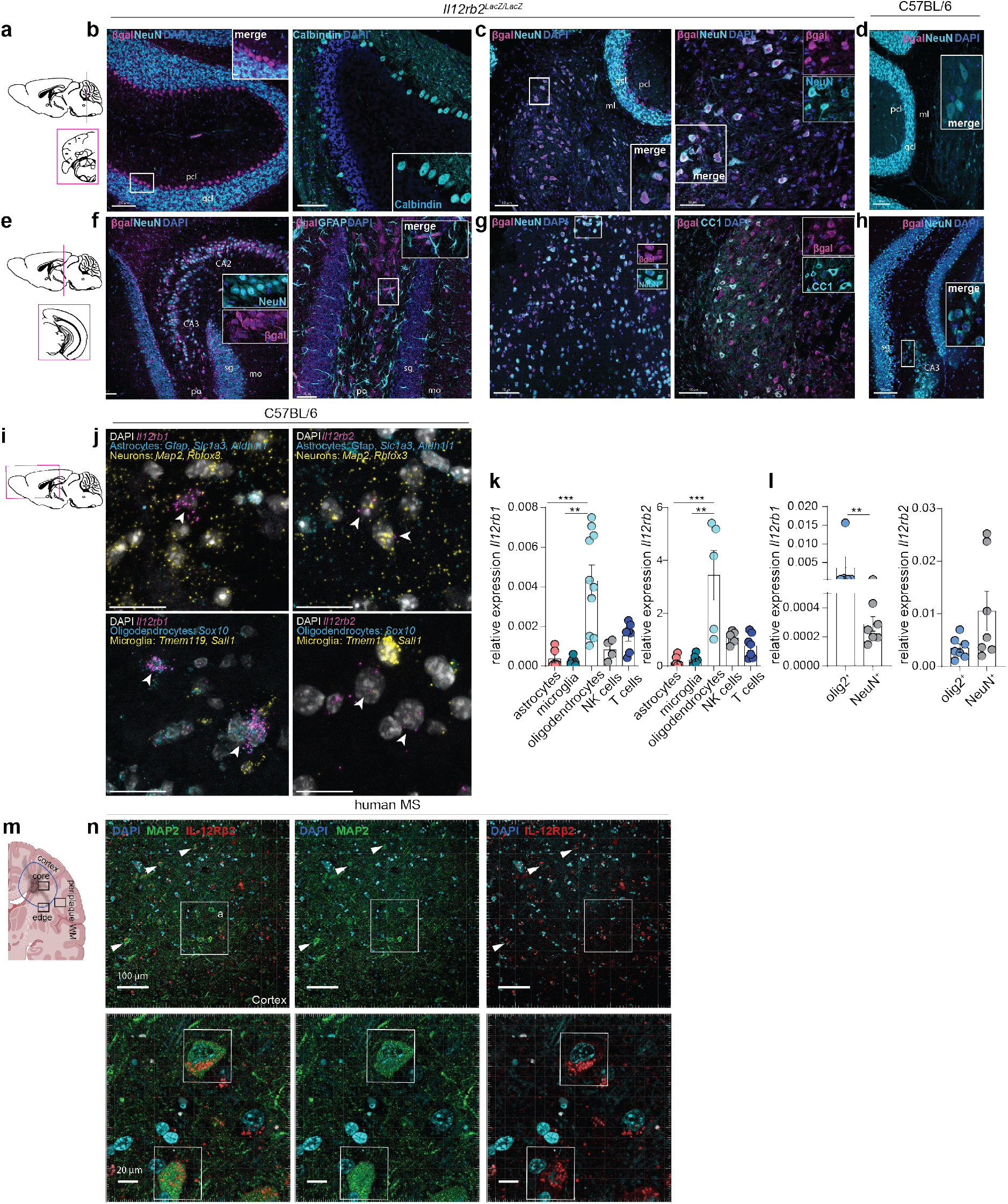
Neurons and oligodendrocytes are molecularly equipped to sense IL-12. (a) Schematic representation of the mouse cerebellum showing region represented by the following images. (b-d) Immunostaining of β-gal^+^Calbindin^+^ and β-gal^+^NeuN^+^ cells in the steady-state cerebellum of *Il12rb2^LacZ^* reporter mice (n≥4mice, ≥5 sections per mouse). Scale bars = 80 μm or 100 μm. (e) Schematic representation of the mouse hippocampus showing region represented by the following images. (f-h) Immunostaining of β-gal^+^NeuN^+^, βgal^-^GFAP^+^ and β-gal^+^CC1^+^ cells in the hippocampal and cortical region of steady-state *Il12rb2^LacZ^* reporter mice (n≥4mice, ≥5 sections per mouse). Scale bars = 80 μm or 100 μm. (i) Schematic illustration of the mouse cerebrum showing region represented by the following images. (j) Multiplexed single-molecule RNA fluorescence in situ hybridization (RNAscope^®^) in steady-state C57BL/6 wt mouse cerebral tissue showing *Il12rb2* gene coexpression in neurons (*Map2, Rbfox3*) and *Il12rb1* coexpression in oligodendrocytes (*Sox10*). Scale bars = 25 μm. (k) *Il12rb1* and *Il12rb2* mRNA expression of astrocytes (CD45^-^CD11b^-^CD31O4^-^ACSA-2^-^, microglia (CD45^low^CD11b^+^CX3CR1^+^), oligodendrocytes (CD45 CD11b CD31 ACSA-2^-^ O4^+^), NK cells and CD3^+^ T cells isolated by FACS from pooled C57BL/6 wt mouse brain and spinal cord tissue at peak EAE (14 dpi). Each symbol represents one mouse (n=4-10). Data shown as mean ± SEM. Kruskal–Wallis test, corrected Dunn’s test. * P < 0.05, ** P < 0.01, *** P < 0.001. (l) *Il12rb1* and *Il12rb2* mRNA expression of oligodendrocyte nuclei (Hoechst^+^Olig2^+^) and neuronal nuclei (Hoechst^+^NeuN^+^) isolated by FACS from the CNS of C57BL/6 wt mice (n=5-7) at peak EAE (14 dpi). Mean ± SEM. Unpaired two-tailed Mann-Whitney test. * P < 0.05, ** P < 0.01. (m) Schematic illustration of human MS brain lesions (created with Biorender.com). (n) Representative immunofluorescence of IL-12Rβ2-expressing MAP2^+^ neurons in brain tissue samples from patients with MS (n=3). Scale bar= 20 μm or 100 μm. GCL, granule cell layer; ML, molecular layer; PCL, Purkinje cell layer; sg (DG-sg), dentate gyrus, granule cell layer; po (DG-po), dentate gyrus, polymorph layer; mo (DG-mo), dentate gyrus, molecular layer.

Similar *Il12rb1/Il12rb2* gene expression patterns were present in the inflamed murine CNS (Extended Data Fig. 2c-f). We further validated the visualization of mRNA transcript or protein expression with qPCR of Fluorescence-Activated Cell Sorter (FACS)-(Fig. 2k) and Fluorescence-Activated Nuclei Sorter (FANS)-purified (Fig. 2l) populations from the inflamed brain. Apart from infiltrating T cells and NK cells, IL-12R subtypes were expressed exclusively on oligodendrocytes and neurons; thus, these cells are the primary CNS-resident cells expressing the IL-12R complex at steady-state and during EAE-induced inflammation.

We next evaluated the presence of the respective IL-12-sensing molecular machinery in the human CNS by examining snRNA-seq datasets consisting of 66,432 nuclei from white matter (WM) lesional tissue of deceased patients with progressive MS and age- and sex-matched controls who had died from non neurological causes^18^ (Extended Data Fig. 2g-i). Analogous to neuroectodermal cell subsets in mouse, human neurons and oligodendrocytes express transcripts encoding IL-12R subtypes (Extended Data Fig. 2h-i). Notably, *IL12RB2* was expressed at a higher level in brain tissue from patients with MS, within all distinct lesion areas and peri-plaque white matter, compared to brain tissue from control patients (Extended Data Fig. 2i). We confirmed neuronal IL-12Rβ2 protein expression in the cortex of MS patients by immunohistochemistry (Fig. 2m-n, Extended Data Fig. 2j).

When we exposed primary mouse neurons (Extended Data Fig. 2k-l) or oligodendrocytes (Extended Data Fig. 2m-n) to recombinant IL-12, we observed a rapid increase of STAT4 phosphorylation-a hallmark of IL-12 receptor signalling. Together, these data demonstrate that in both mice and humans CNS-endogenous neuroectodermal cells are capable of sensing and transducing IL-12 signalling intracellularly.

### IL-12R signalling in the neuroectoderm attenuates neuroinflammation

We then asked whether this ability of the neuroectoderm to sense and respond to IL-12 was functionally linked to IL-12-induced tissue protection in EAE. We crossed *Il12rb2*^fl/fl^ mice to a *Nestin^Cre^* strain^19^ (Fig. 3a), giving progeny in which *Il12rb2* is deleted in the neuroectoderm. We confirmed successful targeting of *Il12rb2* by qPCR of FANS-isolated NeuN^+^ and Olig2^+^ nuclei (Extended Data Fig. 3a), and normal *Il12rb2* expression and function in T- and NK cells by qPCR and IL-12-triggered STAT4 phosphorylation (Extended Data Fig. 3b and c). Strikingly, deleting the IL-12 receptor from the neuroectoderm phenocopied the EAE-hypersusceptibility of the *Il12rb2* full KO strain: *Nestin^Cre/+^Il12rb2^fl/fl^* mice displayed significantly worse EAE clinical scores compared to littermate controls (Fig. 3b, Extended Data Fig. 3d-e).

**Figure 3:**
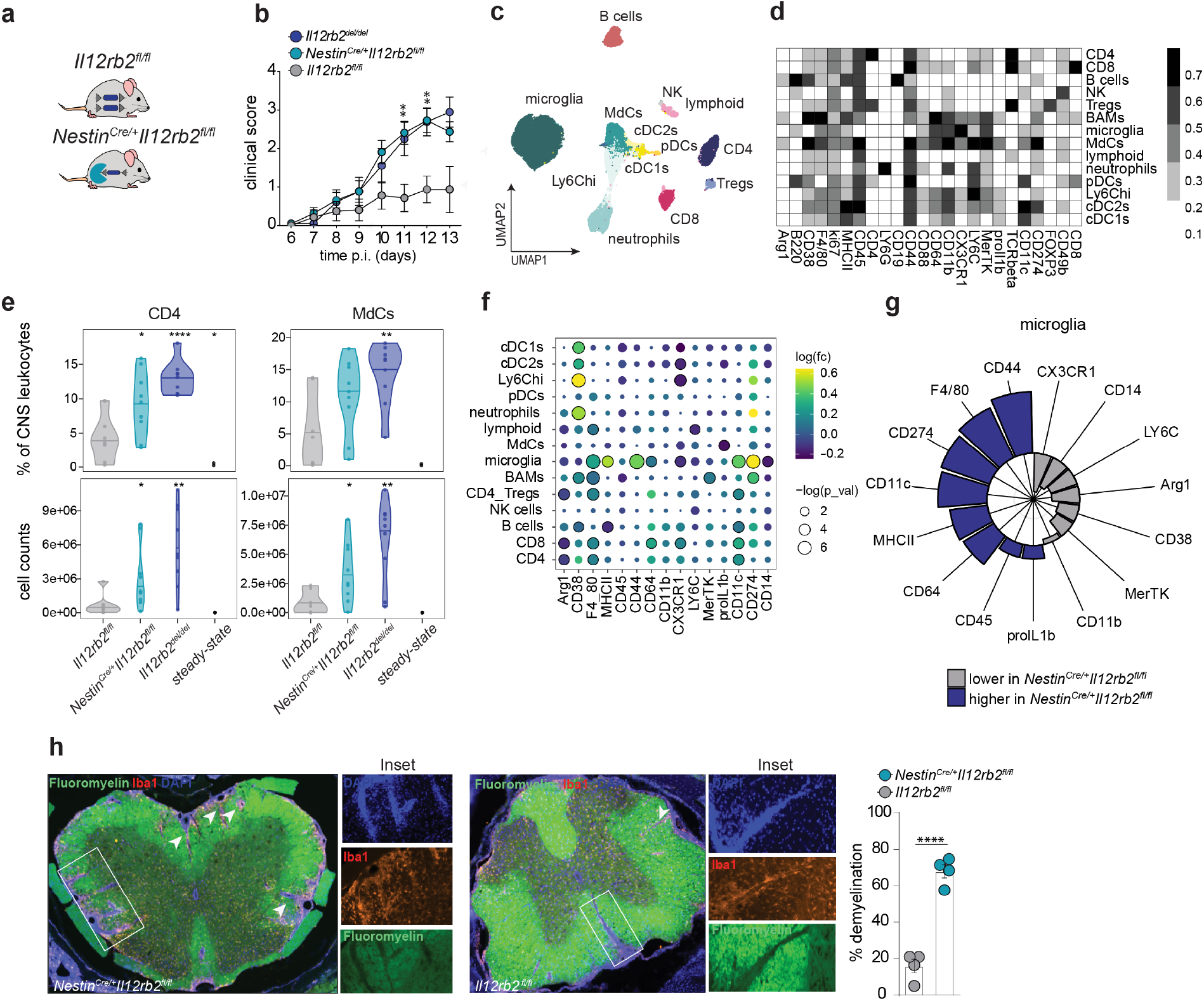
IL-12-sensing in neuroectodermal cells attenuates CNS immunopathology. (a) *Nestin^Cre/+^* mice were crossed to *Il12rb2^fl/fl^* mice to generate *Nestin^Cre/+^Il12rb2^fl/fl^* mice, which lack *Il12rb2* in neuroectodermal cells. (b) Representative clinical disease course of MOG-CFA induced EAE in *Nestin^Cre/+^Il12rb2^fl/fl^* (n=11), littermate controls (n=7) and *Il12rb2^del/del^* (n=9) mice. The disease progress curve depicts pooled data from two independent experiments of similar duration and disease severity (max. EAE score). Data represent mean ± SEM. Two-way ANOVA with Bonferroni post hoc test. * P < 0.05, ** P < 0.01. (c) UMAP of 60,000 cells normalized to sample size and proportional to the absolute cell number per group of *Nestin^Cre/+^Il12rb2^fl/fl^* (n=11), *Il12rb2^fl/fl^* (n=7), *Il12rb2^del/del^* (n=9) at steady-state or at peak EAE (13 dpi). Colours correspond to FlowSOM-guided clustering of cell populations. (d) Median marker expression values for each population. (e) Frequencies (in %) and cell counts of brain-infiltrating CD4^+^ T cells and MdCs at peak disease stage (13 dpi) of cohort (c). Data shown as mean ± SEM; t test. * P < 0.05, ** P < 0.01,*** P < 0.001, ****P < 0.0001. (f) Dot plot depicting expression fold change and p value of the indicated markers per population and (g) effect size for the expression of the indicated activation molecules in microglia isolated at peak EAE (13 dpi) from *Nestin^Cre/+^Il12rb2^fl/fl^* (n=11) and *Il12rb2^fl/fl^* (n=7) mice. (h) Representative white matter immunofluorescence and % myelin loss quantification in coronal spinal cord sections at 19 dpi in *Nestin^Cre/+^Il12rb2^fl/fl^* and control mice (n=4 mice per group). The white matter was stained with FluoroMyelin™ Green dye. Insets highlight active, demyelinating inflammatory lesions with accumulation of infiltrates in their core (DAPI^+^Iba1^+^clusters of phagocytes). Scale bars= 25 μm or 100 μm. Each symbol represents one mouse. Data represent mean ± SEM. Unpaired Student’s t test. * P < 0.05, ** P < 0.01, *** P < 0.001, ****P < 0.0001.

To understand how IL-12 receptor signalling in neuroectodermal cells was linked with inflammatory pathology in EAE, we asked which other cell populations were present in the inflamed CNS during the peak of disease. Using spectral flow cytometry of CNS leukocytes (gated on live CD45^+^ singlets) (Fig. 3c-g, Extended Data Fig. 3f-g), coupled with dimensionality reduction (UMAP^20^) (Fig. 3c) and FlowSOM^21^ clustering (Fig. 3d), we observed significantly higher frequencies of CD4^+^ T cells and MdCs in *Nestin^Cre/+^Il12rb2^fl/fl^* mice with EAE compared to *Il12rb2^fl/fl^* littermates (Fig. 3e, Extended Data Fig. 3g). We also found that microglia in these mice exhibited a proinflammatory signature (Fig. 3f-g), consistent with the exacerbated EAE phenotype of *Nestin^Cre/+^Il12rb2^fl/fl^* mice. Finally, quantitative immunofluorescence confirmed severe inflammation and CNS demyelination in these mice lacking functional IL-12 signalling in neurons and oligodendrocytes (Fig. 3h). Together, our data demonstrate that IL-12 signalling by cells of the neuroectoderm drives immunomodulatory and/or tissue-protective processes which attenuate inflammatory demyelinating CNS pathology in EAE.

### IL-12 prevents early neurodegeneration and sustains trophic factor release in the inflamed CNS

Having identified the critical nature of IL-12 sensing by neuroectodermal cells in EAE, we next sought to precisely disentangle the molecular underpinnings of this yet unperceived neuroimmune crosstalk instructed by IL-12. Importantly, for this analysis, we opted for a time point when IL-12 is already present in the CNS but has not (yet) translated in clinical differences between *Nestin^Cre/+^Il12rb2^fl/fl^* and *Il12rb2^fl/fl^* control mice. Hence, we measured the concentration of IL-12 in brain and spinal cord lysates during EAE induction and disease (Extended Data Fig. 3h-i). We found that IL-12 was barely detectable for almost a week post-immunization, but began to increase from the day of clinical disease onset (day 10 p.i.) (Extended Data Fig. 3h), likely coinciding with the influx of MdCs^22^ (Extended Data Fig. 3j).

Fragile neuroectodermal cells are typically underrepresented in most single-cell omic approaches^23,24^. Instead, we performed snRNA-seq of FANS-isolated Hoechst^+^ nuclei from the cerebellum, brainstem and cervical spinal cord of *Nestin^Cre/+^Il12rb2^fl/fl^* mice and their *Il12rb2^fl/fl^* littermates at the steady-state and onset of clinical symptoms (10 dpi) (Fig. 4a). After quality control and doublet exclusion, snRNA-seq yielded a total of 87,076 single-nucleus transcriptomic profiles, among which 22,018 distinct genes were detected (Extended Data Fig. 4a-b). Unsupervised clustering of these data identified 16 clusters that were assigned to diverse neuronal, glial and other cell types on the basis of known lineage marker genes^15,25–27^ (Fig. 4b, Extended Data Fig. 4c). Among these clusters were 5 populations of neurons reflecting the cellular composition of the cerebellum^15^ [consisting of Granule cells, Purkinje neurons, Golgi cells, Molecular Layer Interneurons (MLI), Unipolar Brush Cells (UBC)] and 3 additional neuronal clusters (excitatory, cholinergic and dopaminergic neurons) classified by neurotransmitter type^26,27^. Non-neuronal clusters consisted of 7 clusters of glial cells [corresponding to oligodendrocytes (Myelin Forming Oligodendrocytes, MFOLs; Mature Oligodendrocytes, MOLs) astrocytes, microglia, oligodendrocyte precursor cells (OPCs) and Bergmann glia] and 1 cluster of vascular and leptomeningeal cells (VLMCs). Amongst all recovered nuclei, 85% were neurons, and 15% were glial cells (Fig. 4c), consistent with previous descriptions^15,28,29^.

**Figure 4:**
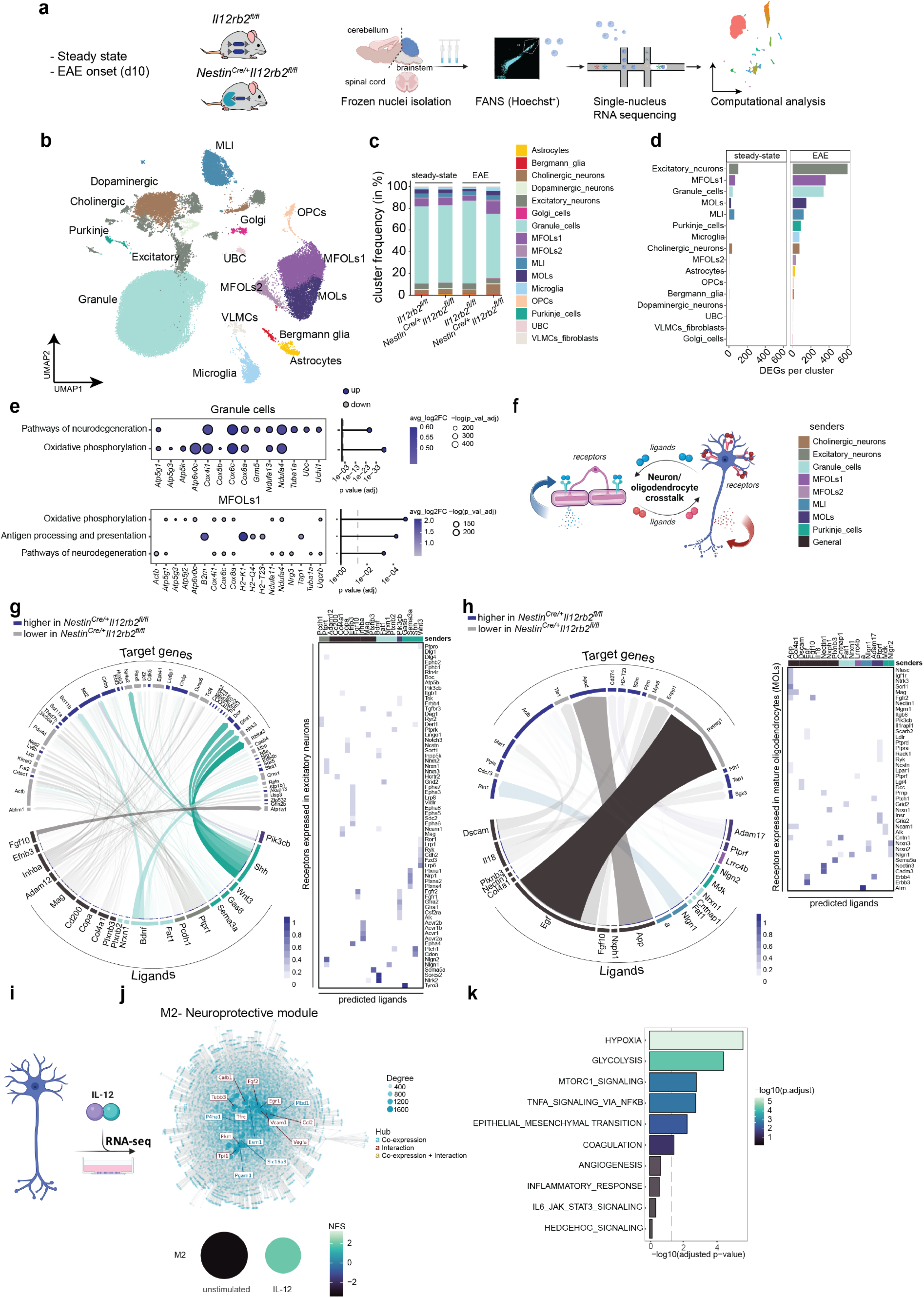
IL-12 signalling promotes neuronal survival, homeostasis and trophic support to oligodendrocytes in the inflamed murine CNS. (a) Overview of the experimental strategy. Illustration was created with Biorender.com. Nuclei were isolated from pooled cerebellum, brainstem (dissected from one brain hemisphere) and cervical spinal cord from *Il12rb2^fl/fl^* and *Nestin^Cre/+^Il12rb2^fl/fl^* male mice (n=1 per genotype, 8-12 weeks old) at the steady-state and early onset of EAE (clinical score<1). Nuclei were lysed, purified by FANS selecting for the Hoechst^+^ fraction (without any doublet exclusion) and used for single-nucleus RNA sequencing. (b) snRNA-seq clustering of 87,076 nuclei by cell type labelled on the basis of known lineage markers and visualized as a uniform manifold approximation and projection (UMAP) plot. Each dot corresponds to a single nucleus and each colour to a cell-type cluster. (c) Percentages of different cell populations split by genotype and disease state. (d) Number of DEGs between *Nestin^Cre/+^Il12rb2^fl/fl^* and *Il12rb2^fl/fl^* mice across the major cell types in steady-state and during EAE. (e) Pathway analysis (KEGG) of DEGs from Granule cells and MFOLs1 of EAE *Nestin^Cre/+^Il12rb2^fl/fl^* versus *Il12rb2^fl/fl^* mice. Top2 or 3 KEGG pathways with g:SCS-corrected p-values < 0.05 are shown with corresponding signature genes. (f) Schematic representation of the receptor-ligand interaction analysis (NicheNet) (created with Biorender.com). Senders were defined by their ability to sense IL-12 (expression of *Il12rb1* and/or *Il12rb2* transcripts) and the magnitude of their differential gene expression (cut-off> 35 DEGs). Both autocrine and paracrine signalling pathways were taken into account. (g) Circos plot (left) showing links between unique ligands from cholinergic neurons, granule cells, Purkinje cells, MLIs, MOLs, MFOLs1 and MFOLs2 (senders; ribbon colour indicates subcluster of origin for each ligand) or general ligands (multiple sender cells; ribbon colour black) and predicted associated DE genes in excitatory neurons (*Nestin^Cre/+^Il12rb2^fl/fl^* versus *Il12rb2^fl/fl^* EAE, blue: higher in *Nestin^Cre/+^Il12rb2^fl/fl^*, grey: lower in *Nestin^Cre/+^Il12rb^fl/fl^* potentially targeted by the ligand-receptor pairs. Transparency indicates interaction strength. Heatmap (right) displaying potential receptors expressed in excitatory neurons associated with each predicted ligand. (h) Circos plot (left) showing links between unique ligands from cholinergic neurons, excitatory neurons, granule cells, Purkinje cells, MLIs, MFOLs1 and MFOLs2 (senders; ribbon colour indicates subcluster of origin for each ligand) or general ligands (multiple sender cells; ribbon colour black) and predicted associated DE genes in MOLs (*Nestin^Cre/+^Il12rb2^fl/fl^* versus *Il12rb2^fl/fl^* EAE, blue: higher in *Nestin^Cre/+^Il12rb2^fl//fl^*, grey: lower in *Nestin^Cre/+^Il12rb2^fl/fl^*) potentially targeted by the ligand-receptor pairs. Heatmap (right) displaying potential receptors expressed in MOLs associated with each predicted ligand. In (g) and (h) the ribbon thickness is proportional to the ligand’s regulatory potential. (i) Primary neuronal cultures were activated with IL-12 (30 ng/μl and 100 ng/μl) for 18 hours and harvested for RNAseq (n=5 replicates per group). (j) Modular differential gene co-expression analysis, depicting module activity across condition and global gene interaction network of module 2 (neuroprotective module) depicting the most connected genes (=hubs). NES, Normalized Enrichment Score. (k) Gene set enrichment analysis depicting the biological components of the neuroprotective module.

When we looked at the distribution of *Il12rb2* mRNA transcripts across the defined cell clusters, we found that it was most abundant in neuronal populations (Extended Data Fig. 4d). *Il12rb1* transcripts were notably enriched within mature oligodendrocyte subsets, specifically in response to immunization. This observation, combined with our evidence of increased demyelination upon loss of IL-12-sensing within the neuroectoderm (Fig. 3h), led us to ask whether IL-12 directly regulated the continuous maturation process of OPCs into mature oligodendrocytes^30^. By examining the top differentially expressed genes (DEGs) and hallmark transcription factors expressed in the oligodendroglial lineage, we noted a differentiation trajectory of oligodendrocyte subsets resembling various cellular states of maturation^31^. However, this maturation process was not affected by the loss of IL-12 signalling (Extended Data Fig. 4e-f) as confirmed by a detailed trajectory inference using RNA velocity (Extended Data Fig. 4g). Hence, the IL-12 mediated tissue-protection likely occurred independently from myelin turnover.

Next, we assessed how the transcriptional profile of the neuroectoderm differed in the presence or absence of IL-12 receptor signalling by interrogating differentially expressed genes (DEGs). This revealed pronounced alterations particularly in excitatory neurons, granule cells, mature oligodendrocytes (MOLs) and myelin forming oligodendrocytes (MFOLs1) suggesting a specific/concentrated action of IL-12 on these populations (Fig. 4d, Supplementary Tables 1-2). In excitatory neurons, we found that several genes implicated in neuronal survival and synaptic plasticity (*Erbb4, Fat2, Neurod1, Camk4, Epha3*) as well as axoglial adhesion molecules required for proper targeting of myelin to axons (i.e. *Cadm3*^32^) were downregulated in *Nestin^Cre/+^Il12rb2^fl/fl^* mice (Extended Data Fig. 4h), indicating that IL-12 counteracts these functional perturbations. Pathway analysis (KEGG) in Granule cells - the most abundant neuronal population - showed enriched expression of genes involved in neurodegeneration (*Cox6c,Cox4i1,Ndufa4,Cox8a,Tuba1a*) and oxidative phosphorylation (*Atp6v0c, Cox6c, Cox4i1, Ndufa4, Cox8a*) in *Nestin^Cre/+^Il12rb2^fl/fl^* mice (Fig. 4e, Extended Data Fig. 4h, Supplementary Table 3) indicating that IL-12 signalling in neurons promotes neuronal survival and homeostasis. Strikingly, AIM2 inflammasome signalling, which protects neurons from genotoxic stress and contributes to homeostatic CNS sculpting^33^, was induced by IL-12 in Granule cells (Extended Data Fig. 4h), thereby decreasing their vulnerability to DNA damage. We observed a similar transcriptomic signature in MFOLs1 and MOLs, where genes involved in cellular responses to oxidative stress (*Ndufa4, Cox6c, Atp6v0c, Atp5g3, Cox4i1*), degeneration (*Ndufa4, Cox6c, Atp5g3, Cox4i1, Tuba1a*) and programmed cell death (*Hif3a, Cst3*) were downregulated by IL-12 (Fig. 4e, Extended Data Fig. 4h, Extended Data Fig. 5a, Supplementary Tables 2-3). In addition, pathways of antigen presentation and processing (*B2m, Tap1, H2-K1, H2-Q4, H2-T23*) (Fig. 4e, Extended Data Fig. 4h, Extended Data Fig. 5a) were upregulated in MOLs and MFOLs1. These features are reminiscent of the recently described “disease-associated oligodendrocyte” signature^34^ and possibly a secondary effect to increased inflammation in *Nestin^Cre/+^Il12rb2^fl/fl^* mice. Together, these findings suggest that the IL-12-mediated neuroectodermal adaptation provides early protection through cell autonomous mechanisms (oxidative stress, cell death) and non-autonomous mechanisms, such as neuronal connectivity and circuit function, to both neurons and oligodendrocytes.

To understand how IL-12-mediated effects on neurons and oligodendrocytes affect intercellular communication and downstream gene expression in the inflamed CNS, we analysed ligand-receptor interactions that are altered upon loss of IL-12 using the NicheNet algorithm^35^ (Fig. 4f-h, Extended Data Fig. 5b-d). We intentionally defined sender cells by their ability to sense and respond to IL-12 (i.e. specific neuronal and oligodendrocyte subpopulations), while recipients included all cells captured by snRNAseq (IL-12R expressing and not expressing). While DEG analysis can only predict direct IL-12 mediated transcriptional changes, this approach allowed us to decipher IL-12 mediated, albeit indirect effects. In total, NicheNet yielded 75 ligand-receptor pairs (Fig. 4f-j, Supplementary Table 4).

Two common patterns of intercellular communication were shaped by IL-12: neuron-intrinsic protection counteracting neurodegeneration and trophic factor support to cells of the neuroectoderm. Specifically, NicheNet revealed IL-12-dependent regulation of receptor-ligand interactions linked to neuronal development, axonal integrity and growth/regeneration, such as BDNF-NTRK2, FAT1-GRID2 and PTPRT-NLGN2 in excitatory neurons (Fig. 4g). Among the most prominent interacting partners we also detected ligand-receptor pairs, such as SHH-PTCH1^36,37^, which most likely represents compensatory efforts aimed at restoring CNS integrity (Fig. 4g). We noted a similar neuroprotective signature emerging in Granule cells, where IL-12 promoted intercellular communication involved in synapse assembly, axon guidance and axon-myelin interactions. These receptor-ligand pairs included NLGN1-NRXN3, NLGN3-CNTNAP2 and NRXN1-NLGN1, which have been implicated in the pathology of MS, neurodegenerative and neuropsychiatric disorders^38,39^ (Extended Data Fig. 5b). Additionally, critical pro-survival cues comprising FGF14-FGFR1/2 and MANF-NPTN, as well as NTIN1-NEO1 engagement were elicited in Granule cells in response to IL-12 (Extended Data Fig. 5b).

The most pronounced theme observed in mature oligodendrocyte populations (i.e. MFOLs1, MFOLs2 and MOLs) affected by the loss of IL-12 sensing was the perturbation of their trophic factor environment. Trophic signals shaping the neuron-oligodendrocyte crosstalk and affected by IL-12 included ADAM17-ERBB4 and neuron-derived NECTIN-FGFR2/3, EGF-ERBB3/ERBB4^40,41^ and FGF10-FGFR2 (Fig. 4h). Neuron-specific ablation of ADAM17, for instance, was shown to impair oligodendrocyte differentiation and myelination *in vivo* and *in vitro*^42^, while EGF has been suggested to enhance remyelination and neurogenesis processes^43^ (Fig. 4h, Extended Data Fig. 5c). A general impact of IL-12 on trophic factor support was also evident in the DEG analysis, where for instance the expression of the Erb-B2 Receptor Tyrosine Kinase 4 (*Erbb4*^44^) and the TAM (Tyro3, Axl, Mer) receptor *Tyro3*^45^, was repressed in *Nestin^Cre/+^Il12rb2*^fl/fl^ mice (Supplementary Table 2). Additionally, the IL-18/IL-18R axis in oligodendrocyte populations, which can interfere with efficient remyelination^46^, was consistently repressed by IL-12 signalling (Fig. 4h, Extended Data Fig. 5c).

Interestingly, many of the interactions affected by IL-12 involved Purkinje cells as senders. Normalizing the number of DEGs to the cluster size revealed that Purkinje cells were in fact strongly influenced by IL-12 signalling (Extended Data Fig. 4i-j). Corroboratively, this neuronal population presented the most pronounced expression of the IL-12 receptor b2 subunit, both at transcript and protein level (Fig. 2b, Extended Data Fig. 4d). By interrogating the top DEGs in this cluster, we observed that genes linked to neurotransmission, synaptic plasticity and synapse assembly (e.g. *Syt1, Nrxn1, Cadm2, Cadm3, and Negr1*) were enriched upon IL-12 sensing in a similar fashion to Granule cells and excitatory neurons. Also here, trophic factors including *Fgf12, Fgf13, Fgf14 and Megf11* were enriched in an IL-12 dependent manner. Taken together, our data suggest IL-12 to be critically involved in neuroprotection and shaping the trophic factor milieu within the inflamed CNS. We observed that the dominant source of trophic factors were neurons (Fig. 4g-h, Extended Data Fig. 5b-c) highlighting their function as critical homeostatic gatekeepers. Conversely, dysregulation or loss of trophic factor release by neurons in the absence of IL-12 may further propagate the degeneration of neurons, oligodendrocytes and possibly other cell types.

Of note, our data do not allow to fully exclude the possibility that transcriptional changes in the neuronal compartment are secondary to the different levels of inflammation observed between *Nestin^Cre/+^Il12rb2^fl/fl^* and *Il12rb2*^fl/fl^ mice. Unfortunately, neuroprotective features cannot be tested at steady-state due to the absence of inflammation, and therefore IL-12, in the homeostatic CNS (Extended Data Fig. 3h). To overcome these limitations and to solidify the findings of the snRNAseq dataset, we performed bulk RNA-sequencing (RNA-seq) of primary murine neurons stimulated with recombinant IL-12 (Fig. 4i-k, Extended Data Fig. 5e-g). Principal component analysis (PCA) revealed a dose-dependent response to IL-12 in the neuronal transcriptome (Extended Data Fig. 5e). Our computational analysis identified 1,878 significantly DEGs (p≤0.01, log ratio ≥0.5) (Supplementary Table 5). Several of the upregulated genes (1091 in total) have been linked to promoting neurogenesis (*Fpr1, Mbdi1*), neuroprotection (*Timp1, Angptl4*) and enhancing remyelination (*Timp1, Matn2*) (Extended Data Fig. 5f). All IL-12-responsive-neurons displayed pronounced *Serpina3n* expression, encoding a Granzyme B (GrB) inhibitor which prevents direct neuronal killing by T cells and NK cells and induces neuroprotection in EAE^47^ (Extended Data Fig. 5f). Confirming our *in vivo* data, IL-12 triggered the secretion of trophic factors for oligodendrocytes including *Lif and Il33* (Extended Data Fig. 5f). Modular gene co-expression network analysis revealed 10 groups of genes (modules) whose expression was jointly altered by IL-12 (Fig. 4j). One of the predicted modules (M2 or “neuroprotective” module), which was induced by IL-12, harboured gene “hubs” (i.e. genes strongly co-expressed and/or interacting with other genes or both) linked to neurogenesis and neuroprotection (*Fgf2, Vegfa, Egr1*), axonal regeneration (*Tubb3*), OPC mobilization (*Cc12*), endogenous myelin repair (*Vcam1, Fgf2*) and neuronal metabolism [glycolysis (*Tpi1*), copper ion transport (*Slc16a3*)] (Fig. 4k). In fact, the neuroprotective module was highly related to glycolytic activity in neurons (Fig. 4k) directly confirming our *in vivo* snRNAseq data where we observed that the loss of IL-12 led to an increase in oxidative phosphorylation across neuroectodermal cell subsets (Fig. 4e). Furthermore, IL-12 directly triggered pathways known to exert pro-survival effects in CNS pathology; for instance hypoxia^48^ and the mTORc1^49^ signalling pathway (Fig. 4k). Collectively, these results confirm and further validate a direct neuroprotective and neurotrophic effect of IL-12.

## Discussion

The discovery of IL-12 coincided with understanding its powerful effects on NK cells and only later on T cells as well^50,51^. IL-12 is firmly linked with type 1 immunity^52^ and is widely held as a major driver of inflammation. Thus, the observation that IL-12-deficient mice develop stronger tissue inflammation across multiple preclinical models of chronic inflammatory diseases (as reviewed in^53,54^) has represented somewhat of a paradox. Cytokines, however, are pleiotropic and specific cytokine receptors are often expressed across different cell types. Cytokine networks not only involve immune cells, but also various cellular entities outside of the hematopoietic compartment^55^. Here, by combining *in vivo* gene targeting with single-cell omics, we demonstrate an unprecedented function of IL-12 in neuroectodermal cells during neuroinflammation. Specifically, we identified two novel CNS-endogenous neuroectodermal sensors of IL-12: neurons and oligodendrocytes. Whereas conditional deletion of *Il12rb2* in cells of hematopoietic origin was dispensable for IL-12-mediated tissue protection, the loss of IL-12-sensing in neuroectodermal cells leads to hyperinflammation during EAE, phenocopying the *Il12rb2* germline mutation^7^.

SnRNA-seq showed pronounced IL-12 driven transcriptional alterations of the neuroectoderm, specifically in granule cells, excitatory neurons, Purkinje cells and mature oligodendrocyte populations. Our analyses revealed that IL-12 instructed neuroprotective cues and trophic factor support within the inflamed CNS. The neuroprotective features of IL-12 were confirmed by next generation sequencing of primary murine neurons exposed to IL-12. This allowed us to uncouple the direct effects of IL-12 from inflammation driven changes in the transcriptome that would result as consequence rather than the cause of differences in disease severity. The directionality agreement of the two independent sequencing datasets validates the mechanistic concordance of our finding that IL-12 directly triggers pathways exerting pro-survival effects on the neuroectoderm. Thereby, IL-12 sensing during disease onset might counteract early neurodegenerative pathways as recently described in human MS^56^. The observation that the IL-12 receptor is expressed in both mouse and human neuroectoderm raises the possibility that IL-12 might trigger a similar reparative mechanism also in the human CNS. Interestingly, treatment with members of the fibroblast growth factor (FGF) family (some of which were consistently upregulated by IL-12) promotes neurogenesis and remyelination in mouse brains^57,58^. In fact, neurons have been demonstrated to be key players in counteracting CNS tissue damage via trophic factor release in the context of diverse brain ailments^59–61^. In contrast, a recent report demonstrated a detrimental role of IL-12 in Alzheimer’s like disease pathology^17^.

The hyperinflamed phenotype of *Nestin^Cre/+^Il12rb2^fl/fl^* mice was characterised by severe CNS tissue damage coupled with markedly increased inflammatory infiltrate. Increased infiltration might be a consequence of overt tissue damage^62^, alternatively immune cells might be actively recruited by brain resident cells. In fact, the concept of the neuroectoderm as an active immunomodulator, rather than passive bystander of inflammation, has emerged in diverse forms of T cell-mediated autoimmune brain pathologies^63–66^. Neurons, in particular, can influence CNS immune responses in both protective^67^, and detrimental ^68,69^ ways. We found that loss of IL-12 sensing led to increased *Cc12* expression in neurons (Extended Data Fig. 5g), thereby likely contributing to the increased inflammatory infiltrate in *Nestin^Cre/+^Il12rb2^fl/fl^* mice. Our bulk RNA-seq data revealed additional examples of IL-12-driven neuroimmune regulation including the induction of protease inhibitors (*Serpina3n, Serpina3i, Serping1 and Timp1*). IL-12 also suppressed the expression of *Mmp12*, whose inhibition in neurons attenuates axonal pathology and degeneration^70^.

In summary, our study resolves a 20-year-old paradox by elucidating the cellular targets and molecular underpinnings of a neuroimmune crosstalk orchestrated by IL-12 in neuroinflammation. The fact that IL-12 can directly initiate neuroprotection may provide a new avenue of localized disease intervention in both neuroinflammatory and degenerative diseases.

## Methods

### Mice

All mice were bred on a pure C57BL/6 background (The Jackson Laboratory, #000664) and were maintained under specific-pathogen free conditions. Knockout-first (promoter-driven) *Il12rb2^fl/fl^* and *Il12rb2^LacZ^* mice were generated as previously described^12^. B6.C-Tg(CMV-Cre)1Cgn/J ^71^ (The Jackson Laboratory, #006054) were crossed to *il12rb2^fl/fl^* mice to generate *CMV^Cre/Cre^Il12rb2^fl/fl^* or *Il12rb2^del/del^* full knock-out (KO) mice. B6.Cg-Tg(Nes-cre)1Kln/J ^19^ (The Jackson Laboratory, #003771) were bred with *Il12rb2^fl/fl^* mice to generate *Nestin^Cre/+^Il12rb2^fl/fl^* mice. For the genotyping of *Nestin^Cre/+^Il12rb2^fl/fl^* mice, blood and tail biopsies were used to distinguish germline deletion from local recombination, as recommended ^72^. *B6.Cg-Tg(Vav1-icre)iKio* ^73^ (The Jackson Laboratory, #008610) were crossed to *il12rb2^fl/fl^* mice to generate *Vav1^Cre/+^Il12rb2^fl/fl^* mice. *B6.Cg-Tg(Cd4-cre)1Cwi/BfluJ* ^74^ (The Jackson Laboratory, #017336) were bred with *il12rb2^fl/fl^* mice to generate *CD4^Cre/+^Il12rb2^fl/fl^* mice. *NKp46^Cre^* mice were kindly provided by E. Vivier and used to generate *Nkp46^Cre/+^Il12rb2^fl/fl^* mice.

All transgenic mouse strains were bred in-house and C57BL/6 wild type mice were purchased from Janvier. Age- and sex-matched (male and female) 8 to 12 week-old mice were used for all experiments. Pups were euthanized between postnatal day 0 (P0) and 4 (P4) for brain harvesting and primary neuronal and oligodendrocyte cultures. All procedures were reviewed and approved by the Swiss Veterinary Office and performed according to institutional and federal guidelines.

To determine the genotype of *Il12rb2^fl/fl^, Il12rb2^LacZ^* and *Il12rb2^del/del^* the following PCR program and primer pairs from KOMP were used: PCR protocol, 94°C (5 min); 10× [94°C (15 s), 65°C (30 s), 72°C (40 s)]; 30× [94°C (15 s), 55°C (30 s), 72°C (40 s)]; 72°C (5 min).

#### Primers for *Il12rb2^fl/fl^ PCR*

CSD-F, 5’-GATTGCCTTAATGAGTAAGAACCTGGF-3’; (forward)

CSD-ttR, 5’-CACAGACATAAAGCAGGTAAACCAACC-3’ (reverse)

Band size for the WT allele is 250 bp; for *il12rb2^fl/fl^* allele 449 bp

#### Primers for *Il12rb2^del/del^ PCR*

CSD-F, 5’-GATTGCCTTAATGAGTAAGAACCTGGF-3’; (forward)

CSD-R, 5’-CATTTGGAGAAAAGAGACAATGTTGG-3’; (reverse)

Band size for the *il12rb2^del/del^* allele is 513 bp; for the *il12rb2^fl/fl^* allele 1374 bp and for the WT allele: 1150 bp

#### Primers for *Il12rb2^LacZ^ PCR*

CSD-lacF, 5’-GATTGCCTTAATGAGTAAGAACCTGGF-3’; (forward)

CSD-R, 5’-CATTTGGAGAAAAGAGACAATGTTGG-3’; (reverse)

Band size for the *il12rb2^LacZ^* allele is 590 bp

#### Exon specific primers for determining targeting efficiency

5’-CTG CGA GAT CTG AGA CC GT-3’

5’-AACA GGCT CTTC CTCT GG TGT-3’

Product length 126 bp

### EAE induction

EAE was induced by injection of 200 μg of myelin oligodendrocyte glycoprotein epitope MOG35-55 (#RP10245, Genscript) emulsified in complete Freund’s adjuvant (CFA) (#231131, BD). All mice received two subcutaneous (s.c.) injections into the flank, each of 100 μl of the freshly prepared MOG-CFA mixture and a single intraperitoneal injection (i.p.) of 50-100 ng pertussis toxin (PT) (List Biological Laboratories, #179A). Of note, most protocols for EAE induction in C57BL/6 mice apply 200 ng of PT. By reducing the concentration of PT, *Il12rb2*^fl/fl^ mice developed a mild disease, whereas *Il12rb2*^del/del^ mice were still hypersusceptible, but within animal welfare guidelines. The mice received a second injection of pertussis toxin at the same concentration 2 days after the initial EAE induction. EAE clinical scores were defined as follows: no detectable signs of EAE: 0; tail limp at distal end: 0.5; entirely limp tail: 1; limp tail and hind limb weakness (occasional grid test positive): 1.5; unilateral partial hind limb paralysis (one leg constantly falls through the grid: 2; bilateral partial hind limb paralysis: 2.5; complete bilateral hind limb paralysis: 3; complete bilateral hind limb paralysis and partial forelimb paralysis: 3.5; complete paralysis of fore and hind limbs (moribund): 4.

### Generation of bone marrow chimeras

Host animals were irradiated with a split dose of 2 × 550 Rad at a 24-hour interval before receiving an intravenous injection (i.v.) of 5 × 10^6^ donor BM cells. The reconstitution period prior to EAE induction was 6 weeks.

### Isolation of cells from the adult mouse CNS

CNS tissues were processed as previously described^75^. Briefly, mice were sacrificed through CO2 asphyxiation and transcardially perfused with ice-cold PBS. Whole brain and spinal cord were isolated, cut into small pieces and digested with 0.4 mg/ml Collagenase IV (#9001-12-1, Sigma-Aldrich) and 0.2 mg/ml deoxyribonuclease I (DNase I) (#E1010, Luzerna) in HBSS (with Ca2^+^ and Mg2^+^) (#14025-050, Gibco) for 40 min at 37°C. The digested tissue was then mechanically dissociated using a 19-gauge needle and filtered through a 100 μm cell strainer (800100, Bioswisstec). CNS single-cell suspensions were further enriched by 30 % Percoll^®^ (P4937, GE Healthcare) gradient centrifugation (1590 g, 30 min, at 4°C, with no brakes).

### Flow Cytometry

Prior to surface labelling, cells were incubated with purified anti-mouse CD16/32 (clone 93, Biolegend) for 10 min on ice to prevent non-specific binding of primary antibodies. Single-cell suspensions were then directly incubated with the primary surface antibody cocktail in PBS for 20 min at 4°C. Following a wash step with PBS (350 g, at 4°C) the cells were then incubated in the secondary surface antibody cocktail (fluorochrome-conjugated streptavidin) for 20 min at 4°C. For intranuclear labelling with Ki67 and Foxp3, cells were fixed and permeabilized utilizing the Foxp3/Transcription Factor Staining Buffer Set (eBioscience) according to the manufacturer’s instructions and subsequently incubated in the intranuclear antibody mixture overnight at 4°C. Anti-mouse fluorochrome-conjugated monoclonal antibodies (mAbs) used in this study were: CD8 (clone 53-6.7), CD274 (clone 10F.9G2 or clone MIH5), CD45R (clone RA3-6B2), CD38 (clone 90), Granzyme B (clone GB11), I-A / I-E (clone M5/114.15.2), Siglec-F (clone E50-2440), NK1.1 (clone PK136), CD45 (clone 30-F11), CD4 (clone GK1.5), Ly6G (clone 1A8), CD19 (clone 1D3), CD44 (clone IM7), CD64 (clone X54-5/7.1), Ki67 (clone B56 or 16A8), F4/80 (clone BM8 or CI:A3-1), CD62L (clone MEL-14), CX3CR1 (clone SA011F11), CD11b (clone M1/70), Ly-6C (clone HK1.4), MerTK (clone DS5MMER), CD103 (clone 2E7), TCR beta chain (clone H57-597), CD11c (clone N418), CD49d (clone R1-2), Foxp3 (clone FJK-16s), CD90.2 (clone 30-H12), TCRγδ (clone GL3), CD88 (clone 20/70), Arginase-1 (clone A1exF5), IL-1 beta Pro-form (clone NJTEN3), CD14 (clone Sa2-B), CD49b (clone DX5).

Sample data were acquired on a 5L Aurora spectral analyser (Cytek Biosciences), FACS Symphony A5 (BD Biosciences) or LSRII Fortessa (BD Biosciences). Flow cytometry data were analyzed using the FlowJo software (version 10.8.0, Tree Star Inc.) and Rstudio (version 4.0.1). All antibodies were purchased from BD Biosciences, eBioscience, AbD Serotec, Biolegend or Miltenyi. Doublet discrimination was performed based on SSC-A/H, FSC-A/H and dead cell exclusion by using a Fixable Viability Kit (Near-IR staining, Biolegend).

### Phosphorylation of STAT4 and pSTAT4 flow cytometry data analysis

Spleens from steady-state mice were pushed through a 70 μm filter and washed once with 1x PBS (350 g, 10 min, at 4°C). Erythrocytes were removed by incubation with RBC lysis buffer (NH4Cl 8.3 mg/mL, KHCO3 1.1 mg/mL, EDTA 0.37 mg/mL) for 5 min on ice. After lysis, samples were washed once in 1x PBS, re-filtered and seeded in 24-well plates (500.000 cells/well) in RPMI medium (supplemented with PenStrep, 10% FCS, L-glut). Cells were cultured with anti-CD3 (5 μg/mL; 3C11; BioXCell) and anti-CD28 (5 μg/mL; 37.51; Biolegend) for 48-72 hours at 37°C, 5% CO2. Cells were then washed in 1x PBS (350 g, 10 min, at 4°C) and stained with a fixable viability dye (Zombie NIR™ Fixable Viability Kit, Biolegend) in PBS for 20 min at 4°C. Following cell viability staining, the cells were resuspended in pre-warmed (37°C) recombinant IL-12-containing (20 ng/ml; PeproTech) medium for 15 min to induce phosphorylation of STAT4. The samples were fixed by directly adding 4% PFA solution (pH 7.4) to a final concentration of 2% (10 min, room temperature). Cells were centrifuged and immediately resuspended in 1 ml of ice-cold methanol for permeabilization (30 min on ice). Subsequently, cells were washed twice with FACS buffer (1% BSA in PBS). Five μl of anti-mouse pSTAT4 (clone 4LURPIE, PE) were added to the cell pellet right before the addition of the surface antibody cocktail. Cells were incubated for 30 min at 4°C, washed and resuspended in 200 μl of FACS buffer for data acquisition.

### Fluorescence-Activated Cell Sorting

Cell sorting was performed using a BD FACS Aria^™^ III. Anti-mouse fluorochrome-conjugated monoclonal antibodies (mAbs) used in this study were: CD45 (clone 30-F11, Pacific Blue, dilution 1:600), CD11b (clone M1/70, BV510, dilution 1:400), CD44 (clone IM7, BV650, dilution 1:400), CX3CR1 (clone SA011F11, BV605, dilution 1:400), LY6C (clone HK1.4, BV711, dilution 1:400), LY6G (clone 1A8, FITC, dilution 1:400), NK1.1 (clone PK136, PE-Cy7, dilution 1:400), O4 (clone REA576, PE, dilution 1:25), ACSA-2 (REA969, APC, dilution 1:25), CD88 (clone 20/70, Biotin, dilution 1:200), F4/80 (clone CI:A3-1, AF647, dilution 1:200), CD140a (clone APA5, PE-Cy7, dilution 1:400), TER119 (clone Ter-119, APC, dilution 1:200), CD3 (clone 17A2, APC, dilution 1:200), CD31 (clone MEC13.3, PE, dilution 1:200), CD31 (clone 390, BV605, dilution 1:400), CD4 (RM4-5, BV605, dilution 1:400), Streptavidin (PE-Cy7-conjugated, dilution 1:400).

All antibodies were purchased from BD Biosciences, eBioscience, AbD Serotec, Biolegend or Miltenyi. Doublet exclusion was performed based on SSC-A/H, FSC-A/H and dead cell exclusion by using a Fixable Viability Kit (Near-IR staining, dilution 1:500, Biolegend).

### Quantitative RT-PCR (qRT-PCR)

Total RNA was isolated from sorted cells using either the RNeasy Micro Kit (#74004, Qiagen) or the QuickRNA Microprep Kit (#R1051, Zymo Research) according to the manufacturer’s instructions. Complementary DNA (cDNA) was synthesized using the M-MLV Reverse Transcriptase (#28025013, Invitrogen) and qRT-PCR was performed on a CFX384 Touch Real-Time PCR Detection System (Bio-Rad) using SYBR Green (Bio-Rad). Total RNA was isolated from sorted Hoechst^+^ Olig2^+^ and Hoechst^+^ NeuN^+^ nuclei using the RNeasy Micro Kit (#74004, Qiagen). Nuclear RNA was reverse-transcribed to cDNA with the RevertAid H Minus First Strand cDNA Synthesis Kit (Thermo Fisher Scientific) according to the manufacturer’s instructions. Gene expression was calculated as 2^-ΔCt^ relative to *Pol2* as the endogenous control as described in Livak et al. ^76^.

#### Primers for *Il12rb2*

5’-CTGCGAGATCTGAGACCGT-3’; (forward)

5’-AACAGGCTCTTCCTCTGGTGT-3’ (reverse)

#### Primers for *Il12a*

5’-TACTAGAGAGACTTCTTCCACAACAAGAG-3’; (forward)

5’-TCTGGTACATCTTCAAGTCCTCATAGA-3’ (reverse)

#### Primers for *Il12b*

5’-GACCATCACTGTCAAAGAGTTTCTAGAT-3’; (forward)

5’-AGGAAAGTCTTGTTTTTGAAATTTTTTAA-3’ (reverse)

#### Primers for *Il12rb1*

5’-GGACCAGCAAACACATCACCTT-3’; (forward)

5’-GTGATGGCTGCTGCGTTG-3’ (reverse)

#### Primers for *Pol2*

5’-CTG GTC CTT CGA ATC CGC ATC-3’; (forward)

5’-GCT CGA TAC CCT GCA GGG TCA-3’ (reverse)

### Primary neuronal cultures

Dissociated cerebellar cells were prepared and maintained as previously described ^77^. Briefly, cerebellar cells from postnatal day 0-4 (P0-P4) mouse pups (C57BL/6) were dissociated and plated on poly-L-lysine-coated, 12-well plates (90.000 cells per well). Cells were maintained in Neurobasal Plus medium supplemented with B27 Plus (#A3653401, Thermo Fisher Scientific) and 2 mM GlutaMax™ (#35050061, Thermo Fisher Scientific).

### Primary oligodendrocyte cultures

For murine primary oligodendrocytes cultures, brains were collected from P2-P5 pups and dissociated using the Neural Tissue Dissociation Kit (P) (Miltenyi Biotec, 130-092-628) on a gentleMACS Octo Dissociator (Miltenyi Biotec, 130-096-427). The resulting single-cell suspension was labeled with O4-conjugated magnetic MicroBeads (Miltenyi Biotec, 130-096-670) and passed through LS columns (Miltenyi Biotec, 130-042-401) to positively select for oligodendrocytes. After isolation, the primary oligodendrocytes were plated at a density of 100,000 cells per 24 well plate on 10% PLL (Sigma, P4832-50ML) and 1.5% laminin (Sigma, L2020) coated plates (Corning, 3524) in 500 μl growth medium (MACS Neuro Medium (# 130-093-570) containing 2% MACS Neuro Brew-21 (# 130-093-566), 1% penicillin/streptomycin (Gibco, 15070063) and 0.5 mM L-Glutamine (25030081), 10 ng/mL Human PDGF-AA (# 130-093-977), and 10 ng/mL Human FGF-2 (# 130-093-837). At days two and four of culture, 250 μl of medium were removed and an additional 250 μl of fresh growth medium were added. At day four, cells were stimulated as indicated with 100 ng/μl of IL-12p70 (Preprotech, 210-12) and harvested for Western Blot analysis.

### Western Blot

Primary neuronal cells (DIV14) or oligodendrocytes were stimulated with 100 ng/μl of recombinant murine IL-12p70 (#210-12, Peprotech) for 5 min, 10 min or 15 min. After stimulation, neurons and oligodendrocytes were harvested and lysed in EBC buffer [50 mM Tris (pH 8.0), 120 mM NaCl, 0.5% Nonidet P-40] containing complete miniprotease inhibitor (#11836153001, Roche Diagnostics) and phosphatase inhibitor mixture 1 and 2 (#P2850 and #P0044, Sigma-Aldrich) for 20 min at 4°C. The clear cell lysate was collected by centrifugation (10,000 g, 4 °C, 20 min) and supernatants were stored at −80°C. Samples were run on Tris-glycine polyacrylamide gels and proteins were transferred to PVDF membranes. Primary antibodies (Cell Signaling: Stat4 #2653; Phospho-Stat4-Tyr693 #4134) were incubated with the membranes in Tris-buffered saline with 0.05% Tween 20 (TBST), including 5% Western Blocking Reagent Solution (#11921673001, Roche) overnight at 4°C. Membranes were washed five times for 5 min in TBST before the addition of secondary antibodies. HRP-coupled donkey secondary antibodies (1:30,000) and fluorochrome-coupled donkey secondary antibodies (1:20,000) were incubated for 30 min at room temperature, and membranes were washed again five times for 5 min in TBST. Fluorescent signals were captured using the Odyssey^®^ CLx Imager. SuperSignal West Pico Chemiluminescent Substrate (#34579, Thermo Fisher Scientific) was applied to visualize HRP-labeled antibodies and developed using the FUJIFILM Luminescent Image Analyzer LAS-1000 plus & Intelligent Dark Box II (Fujifilm). Western blot membranes were re-probed for p-Stat4 and Stat4 using a mild stripping protocol from Abcam: briefly, membranes were incubated for 5-10 min with mild stripping buffer (200mM gylcine, 20mM SDS, 0.01% Tween 20, pH 2.2) followed by 10 min incubation with PBS and 5 min incubation with TBST. This process was repeated twice. Efficiency of stripping was assessed by chemiluminescent detection. When stripping was judged satisfactory, the membranes were rinsed and incubated with primary antibody. Images were processed and analyzed using ImageJ ^78^.

### Multiplex RNA-fluorescence in situ hybridization

Frozen brain tissue was placed in a tissue mould (#SA62534-15, Sakura) and submerged in Tissue-Tek^®^ freezing medium (#4583, Sakura). 10 μm-thick brain tissue sagittal sections were cut using a cryostat (Thermo Fisher Scientific, HM 560), mounted on SuperFrost Plus slides (#500621, R. Langenbrink) and dried for 1 hour at −20 °C. Tissue processing for RNAscope^®^ multiplex staining was carried out following the manufacturer’s protocol for fresh-frozen sections. Briefly, tissue was fixed in freshly prepared 4 % PFA (pH 7.4) for 30 min at 4 °C, followed by alcohol dehydration and exposure to H2O2 for 10 min then Protease IV (#322340, Bio-Techne) for 30 min, both at room temperature. Sections were then incubated for 2 hours with target probes at 40 °C in a HybEZ™ Hybridisation System (#321711, Bio-Techne). The following RNAscope^®^ target probes were used: Mm-Il12rb1 (#488761, Bio-Techne), Mm-Il12rb2 (#451301, Bio-Techne), Mm-Aldh1l1-C2 (#405891, Bio-Techne), Mm-Slc1a3-C2 (#430781, Bio-Techne), Mm-Gfap-C2 (#313211, Bio-Techne), Mm-Sox10-C2 (#435931, Bio-Techne), Mm-Tmem119-C3 (#472901, Bio-Techne), Mm-Sall1-C3 (#469661, Bio-Techne), Mm-Rbfox3-C3 (#313311, Bio-Techne), Mm-Map2-C3 (#431151, Bio-Techne). Signal amplification was achieved using the RNAscope^®^ Multiplex Fluorescent Kit v2 (#323110, Bio-Techne), following the manufacturer’s protocol. Probes were labelled with Opal™ 520 (1:500, C2 probe, FP1487001KT, Perkin Elmer), Opal™ 570 (1:500, C1 probe, FP1488001KT, Perkin Elmer) and Opal™ 690 (1:500, C3 probe, FP1497001KT, Perkin Elmer) and three-dimensional image stacks (1 μm step size, 40x objective) of stained sections were acquired on a Leica TCS SP5 confocal laser scanning microscope using a HCX PL APO lambda blue 63× oil UV objective controlled by LAS AF scan software (Leica Microsystems).

### CNS tissue homogenization and protein extraction

Murine brain and spinal cord were isolated as described above and snap-frozen in liquid nitrogen for storage at −80°C. For protein extraction, samples were subjected to mechanical homogenization in Cell Lysate Buffer (1mM EGTA, 1 mM EDTA, 1% Triton X-100, Tris-buffered saline (TBS) buffer pH 7.5 (20 mM Tris, 150 mM NaCl) filled up to 30 ml with ddH2O). All buffers were kept on ice and the final Cell Lysate Buffer was supplemented with cOmplete™ Mini EDTA-free Protease Inhibitor Cocktail (#11836170001, Roche, 1 tablet per 25 ml) and PhosSTOP™ (#4906845001, Roche, 1 tablet per 10 ml) immediately before use. The initial homogenization was performed mechanically (2 metal beads/sample) using a TissueLyserLT (#85600, Qiagen) homogenizer (50 oscillations/s for 10 min), followed by subsequent incubation on ice for 30 min and centrifugation (full speed, 10 min, 4°C). Protein concentrations of each sample were determined using the Pierce™ BCA Protein Assay Kit (#23225, Thermo Fisher Scientific) according to the manufacturer’s protocol using the Photometer Tecan Infinite^®^200M (Tecan).

### ELISA analysis

ELISA to detect IL-12/IL-23 total p40 (#DY499, R&D) was performed according to the manufacturer’s instructions. All samples were analyzed in duplicate. Absorption was read at 450 and 570 nm (for wavelength correction) on a microplate reader (Infinite 200M, Tecan) and analyzed using the Magellan Software (Tecan).

### Immunofluorescence and imaging of murine brain tissue

Mice were euthanized through CO2 asphyxiation and transcardially perfused with ice-cold PBS. Brains were carefully removed, fixed in 4% PFA (# 11762.00500, Morphisto) for 24 hours at 4°C, rinsed in PBS and then incubated in 30% sucrose in PBS at 4°C for 24-72 hours. Each brain hemisphere was embedded sagittally or coronally in Cryo Embedding Medium (# 41-3011-00, Medite) and frozen on dry ice. Free-floating brain sections were cut at a 35-45 μm thickness using a Hyrax C60 cryostat (Zeiss). Sections were washed three times with PBS and incubated in 10% normal goat serum (#PCN5000, ThermoFisher Scientific) and 0.5% Triton X-100 (Sigma-Aldrich, cat # T8787-100ML) in PBS (2 hours, room temperature) for blocking and permeabilization. Sections that were labelled with antibodies derived from mouse were additionally blocked for another 45 min at room temperature with the M.O.M kit Mouse Ig Blocking Reagent (90ul in 2.5ml PBS; # BMK-2202, Vector Laboratories). Sections were incubated with the primary antibody cocktail overnight at 4°C in 5% normal goat serum and 0.5% Triton X-100 in PBS. The following primary antibodies were used: rat anti-GFAP (#13-0300, Thermo Fisher Scientific; 1:400), rabbit anti-Calbindin (#Abcam; 1:500), mouse anti-NeuN (#MAB377, Chemikon 1:500), mouse CC1 Anti-APC (Ab-7) (#OP80-100UG, Merck) and mouse anti-Olig2 (#66513-1-IG, Thermo Fisher Scientific; 1:200). After washing, sections were with the respective fluorochrome-conjugated secondary antibodies (goat anti-mouse/anti-rabbit/anti-rat, streptavidin etc, conjugated to AlexaFluor 488, 594, 633, 647, all purchased from ThermoFisher Scientific; 1:500) for 2 hours at room temperature in 5% normal goat serum and 0.5% Triton X-100 in PBS. After labelling, sections were mounted in SlowFade Gold antifade reagent with 4’, 6-diamidino-2-phenylindole (DAPI) (#P36931, Thermo Fisher Scientific) for nuclei counterstaing. For co-labelling with rabbit-anti-beta galactosidase (#559761, MP Biomedicals; 1:5000 or former Cappel codice 559762 (Rabbit IgG)), free-floating brain sections were blocked and labelled with the respective primary and secondary antibodies in FBT blocking buffer (5% fetal bovine serum, 1% bovine serum albumin, 0.05% Tween 20, 10 mM Tris-HCl pH 7.5, 100 mM MgCl2 in H_2_O), as described before^79^. Imaging was performed on a STELLARIS 5 (Leica) microscope with a 20X multi-immersion objective with 1024×1024 pixels. High-resolution images were acquired in frames with a line average of 16 or 32. Imaris imaging software (Bitplane, Zurich, Switzerland) was used for image processing and merging.

### Immunostaining of human brain tissue

Brain tissue from three patients with MS was obtained from the archives of the Institute of Neuropathology at the University Hospital Zurich, Switzerland. Informed consent for autopsy was given by the next-of-kin in all cases. Case series do not need institutional review board approval according to Swiss legislation. Immunohistochemistry was performed on formalin-fixed paraffin-embedded (FFPE) tissue sections (2-3 μm thick) as follows: for epitope demasking, deparaffinised sections were heated in a microwave histoprocessor (HistosPro) with DAKO Target Retrieval solution (citrate buffer pH 6.0 (#S2367, DAKO) or pH 9.0 (#S2367, DAKO)), washed with distilled water and transferred to 0.3% H2O2 for 10 min to block endogenous peroxidase. The sections were then washed with PBS and subsequently incubated with blocking buffer (5% NGS/PBS+Triton 0.1%) for at least 20 min. Thereafter, the sections were prepared for primary and secondary antibody incubation. The following primary antibodies were used (diluted in blocking solution): rabbit anti-IL-12Rβ2 (#NBP1-85983, Novus Biologicals; 0.7μg/ml), rabbit IgG purified (#PP64-10 KC, Merck; 0.7μg/ml) and anti-NeuN (#MAB377, Chemikon 1:100). Antibody binding was visualized using Dako EnVision + Dual Link System-HRP (DAB+) staining (DAKO) according to the manufacturer’s procedure. We recorded digital images of tissue sections using an Olympus BX41 light microscope with an Olympus ColorView IIIu camera and Olympus Cell B image acquisition software. For Immunofluorescence stainings the endogenous enzyme blocking step with 0.3% H2O2 was omitted, and rabbit anti-IL-12Rβ2 (#NBP1-85983, Novus Biologicals, 1:1000) was combined with mouse anti-Map2 (#M4403, Sigma, clone HM-2, 1:100) to co-label neurons. The secondary antibody mixture contained anti-rabbit AlexaFluor-555 and anti-mouse AlexaFluor-488 (all purchased from ThermoFisher Scientific; 1:250). Nuclear counterstaining was performed using SlowFade Gold antifade reagent with DAPI (Invitrogen). Imaging was performed on a STELLARIS 5 (Leica) microscope with a 20X multi-immersion objective. Imaris imaging software (Bitplane, Zurich, Switzerland) was used for image processing.

### Quantitative histopathological analysis of spinal cords from mice with EAE

Mice were sacrificed through CO2 inhalation and transcardially perfused with ice-cold PBS. Spinal columns were carefully dissected out and fixed in 4% PFA (# 11762.00500, Morphisto) for 24 hours at 4°C, rinsed in PBS followed by decalcification in 0.5 M, pH=8 EDTA solution (#A3145,0500, Axonlab) at 4°C for 5 consecutive days while rotating. The spinal cords were then dissected into 3 parts (cervical, thoracic and lumbar) and cryoprotected in 30% sucrose in PBS at 4°C for 24-72 hours. Each spinal cord fraction was embedded coronally in Cryo Embedding Medium (# 41-3011-00, Medite) and frozen on dry ice. Frozen sections were cut at a 20-25 μm thickness using a Hyrax C60 cryostat (Zeiss). Subsequent blocking, staining steps and mounting were performed as described above. For the quantification of demyelination and inflammation in EAE, sequential 200 μm-apart spinal cord sections were fluorescently labeled with rabbit anti-Iba1 (#019-19741, Wako; 1:200) and FluoroMyelin™ Green Fluorescent Myelin Stain (#F34651, Thermo Fisher Scientific). For automated imaging, fluorochrome-labeled sections were acquired on a Zeiss Axio Scan Slidescanner using a 20X objective lens. The following fluorescence filters were used: AF405, AF488, AF546 or AF594 and AF647. The semi-automated quantification was performed on 20-30 images of n=3-4 mice/group. Acquired images were cropped in Fiji in order to remove all pixels outside of the spinal cord, and cropped images were then processed in Ilastik^80^. Ilastik was trained to recognize lesions (DAPI bright, fluoromyelin low), white matter (DAPI low, fluoromyelin high), gray matter (DAPI low, fluoromyelin low) and background (no signal), and to calculate the probability that each pixel belonged to each category. These probabilities were exported to Fiji as 8-bit images and thresholded using the default algorithm. The areas of lesions and white matter were then measured, and the percent demyelination was calculated as follows: 100*area of lesions/ (area of lesions+ area of white matter).

### Bulk RNA isolation of primary neuronal cultures

Dissociated cerebellar cells were prepared and maintained as described above and previously^77^. At DIV14, primary neuronal cells were stimulated with 30 ng/μl or 100 ng/μl of Recombinant Murine IL-12p70 (#210-12, Peprotech) for 18 hours. Cerebellar neurons were harvested and lysed in 300 μl RLT buffer of the RNeasy Micro Kit (#74004, Qiagen) and total RNA was extracted according to the manufacturers’ protocol.

### Library preparation, cluster generation and next-generation sequencing

The quality of the isolated RNA was determined with a Fragment Analyzer (Agilent, Santa Clara, California, USA). The TruSeq Stranded mRNA (Illumina, Inc, California, USA) was used in the succeeding steps. Briefly, total RNA samples (100-1000 ng) were poly A enriched and then reverse-transcribed into double-stranded cDNA. The cDNA samples were fragmented, end-repaired and adenylated before ligation of TruSeq adapters containing unique dual indices (UDI) for multiplexing. Fragments containing TruSeq adapters on both ends were selectively enriched with PCR. The quality and quantity of the enriched libraries were validated using the Fragment Analyzer (Agilent, Santa Clara, California, USA). The product is a smear with an average fragment size of approximately 260 bp. The libraries were normalized to 10nM in Tris-Cl 10 mM, pH8.5 with 0.1% Tween 20. The Novaseq 6000 (Illumina, Inc, California, USA) was used for cluster generation and sequencing according to standard protocol^81^. Sequencing configuration was single-end 100 bp.

### Bulk transcriptome analysis of primary neuronal cultures

The raw reads were aligned to mouse genome build GRCm39 using the STAR aligner, with FeatureCounts used to calculate read counts per gene based on GENCODE gene annotation release M26.

Bulk RNA-seq analysis was performed using the edgeR and DEseq2 packages^82,83^. Counts were counts-per-million (CPM) normalized, log2 transformed and only genes were retained if expressed at one CPM in at least two samples. PCA was computed using the BiocGenerics package after variance stabilizing transformation and visualized using ggplot2. Differentially expressed genes between conditions were computed using gene-wise negative binomial generalized linear models and applying a Benjamini-Hochberg correction. Heatmaps of DEGs were generated using pheatmap.

Differential analysis of modular gene co-expression was performed using the R implementation of CEMiTool usind default parameters^84^. For gene set enrichment analysis and interactome integration murine gene names have been converted to human orthologs using biomaRt. Gene set enrichment analysis of gene modules was performed using the hallmark gene set collection from MSigDB^85^.

Interactome data from the STRING database version 11 was integrated after filtering on interactions with a minimum score of 200^86^ to obtain gene co-expression and interaction networks for each module.

### Single nuclei preparation and Fluorescence activated nuclei sorting

Single nuclei suspensions were prepared as described before^17^. Mouse cerebellum and brainstem (of one brain hemisphere) and cervical spinal cord (C1-C2) were harvested from adult male mice (8-12 weeks old) and immediately snap-frozen in liquid nitrogen and stored at −80°C until further processing. Nuclei were isolated with the EZ PREP lysis buffer (#NUC-101, Sigma). Tissue samples were homogenized using a glass dounce tissue grinder (#D8938, Sigma) (25 strokes with pastel A, 25 strokes with pastel B) in 2 ml of ice-cold EZ prep lysis buffer and incubated on ice with an additional 2 ml of ice-cold EZ PREP lysis buffer. During incubation, 1 μM of Hoechst (#H3570, Thermo Fisher Scientific) dye and 40 U/μl of RiboLock inhibitors (#EO0382, Thermo Fisher Scientific) were added to the homogenate. Following incubation, the homogenate was filtered through a 30 μM FACS tube filter. Nuclei were sorted based on the fluorescent Hoechst signal using a BD FACS Aria sorter III with an 85 μm nozzle configuration at 4°C, directly into PBS + 4% BSA + RiboLock inhibitors (40 U/μl) (Cat # EO0382). As CNS nuclei vary strongly in size, no doublet exclusion was performed based on FSC or SSC to avoid bias against nucleus size. Nuclei were then counted based on brightfield image and Trypan Blue staining (Cat # 15250061) using a Neubauer counting chamber and a Keyence BZX-710 microscope.

### Droplet based single-nucleus RNA sequencing

Immediately after extraction, 17.500 sorted nuclei per sample were loaded onto a Chromium Single Cell 3’ Chip (10X Genomics) and processed for the single-nucleus cDNA library preparation (Chromium Next GEM Single Cell 3’ Reagent Kits v3.1 protocol). 50.000 reads per nucleus were sequenced using the Illumina Novaseq 6000 #1 platform according to the manufacturer’s instructions without modifications (R1= 28, i7= 10, i5= 10, R2=90). Preparation of cDNA libraries and sequencing were performed at the Functional Genomics Center Zurich. CellRanger software (v6.0.2) was implemented for library demultiplexing, barcode processing, fastq file generation, gene alignment to the mouse genome (GENCODE reference build GRCm39), and unique molecular identifier (UMI) counts. We implemented the “include-introns” option for counting intronic reads, as the snRNA-seq assay captures unspliced pre-mRNA as well as mature mRNA. For each sample, a CellRanger report was obtained with all the available information regarding sequencing and mapping parameters. All samples were merged into a matrix using CellRanger (cellranger -aggr function).

### Quality control and data pre-processing

Starting from the filtered gene-cell count matrix produced by CellRanger’s in-built cell calling algorithms, we proceeded with the SCANPY workflow in Python^87^. Quality control (QC) processing, graph-based clustering, visualizations and differential gene expression analyses for cluster assignment were performed using SCANPY (v1.8.2.). In brief, we filtered out potentially low-quality nuclei as follows; we exclusively included genes which were expressed in at least 3 nuclei; we also selected nuclei with more than 1000 total counts per nucleus, more than 500 genes expressed in the count matrix and less than 5% of counts in mitochondrial genes. For doublet discrimination we implemented the Scrublet^88^ package with default parameters. Lastly, we corrected for batch effect across datasets using a mutual Nearest neighbour method (MNN).

### Dimensionality reduction, clustering visualization and cell type identification

All remaining variable genes were used for downstream analyses. We normalized and log-transformed UMI counts to a depth of 10,000. We found variable genes using “FindVariableFeatures” with min_mean=0.0125, max_mean=3, min_disp=0.5. In addition, we used the regression algorithm to remove any unwanted sources of variation from the dataset, such as cell cycle, sequencing depth and percent mitochondria. Before performing PCA, each gene was scaled applying Z-score normalization. Dimensionality reduction was performed using the PCA and UMAP algorithms. Next, we proceeded to clustering using the Leiden graph-clustering method. The clustering algorithm was applied with a resolution of 0.8 and identified in 32 initial clusters. For assigning clusters to cell types, we defined a list of marker genes per cluster and ranked genes applying the Wilcoxon Rank-Sum test. The assignment of cell type identity to clusters was based on known linage markers in line with previously published snRNAseq^15^ studies and atlases^26,27^.

### RNA velocity

Annotations of unspliced/spliced reads were obtained using velocyto CLI with default parameters. The number of analyzed reads of the manually assigned oligodendrocyte and OPC clusters were normalised per group and genotype. In brief, we merged unspliced counts with the preprocessed, normalized, scaled and annotated spliced count matrix via the scvelo.utils.merge function and proceeded with the RNA velocity analysis using the velocyto (0.17) and scVelo (v0.2.4) workflow^89^. Before running the dynamical model, we computed moments for velocity estimation implementing the values n_pcs = 20 and n_neighbors = 30. We ran the dynamical model to learn the full transcriptional dynamics of splicing kinetics, transcriptional state and cell-internal latent time across the complete dataset.

### Differential gene expression and pathway analysis

For differential expression between *Il12rb2*^fl/fl^ and *Nestin^Cre/+^Il12rb2^fl/fl^* mice genes were prefiltered for genes that are expressed at least 10% in the respective cluster in one of the groups and demonstrate a more than 0.25 log2FC enrichment between groups. Differentially expressed genes were identified using a non-parametric Wilcoxon Rank Sum test and Bonferroni correction was applied to correct for multiple testing. Pathway analysis was performed using the gprofiler2 package. Differentially expressed genes were sorted based on adjusted p-value and an ordered query using pathways of the KEGG database was submitted. Resulting p-values were corrected for multiple testing using the g_SCS implementation in gprofiler2. Bar graphs, volcano plot, dot plots and lollipop plots were drawn in ggplot2.

### Analysis of ligand-receptor interactions

Ligand-receptor interaction analysis has been carried out using NicheNet and default parameters^35^.Sender cells were defined based on their ability to express *Il12rb2* and demonstrate transcriptional changes after genetic deletion using *Nestin^Cre/+^Il12rb2^fl/fl^* mice (MOLs, MFOLs1, MFOLs2, Purkinje cells, Excitatory neurons, Cholinergic neurons, Granule cells and MLI). For each receiver cell NicheNet predicted the ligands that best explain differentially expressed genes upon *Il12rb2* deletion. Top predicted ligands and the corresponding target genes for each receiver cell type were visualized in circus plots using the circlize package. Dot plots showing the respective ligand expression across sender cells were generated in ggplot2. Heatmap of top predicted ligands in sender cells and corresponding receptors in recipients were generated using the pheatmap package.

### Re-analysis of single-nucleus RNA-seq of MS tissue

Publicly available gene expression matrix from Absinta *et* al. (accessed at GSE180759) ^18^ was re-analyzed for the expression of *IL12RB1* and *IL12RB2* using the Seurat v3 R-based pipeline^90^.

### Fluorescence Activated Nuclei Sorting for bulk RNA isolation

Nuclei of murine brains and spinal cords were isolated and sorted as described above. For specific labeling of neuronal and oligodendrocyte nuclei, the homogenates were directly stained with Hoechst, NeuN (#ab190195, Abcam; 1:200) and Olig2 (#ab225100, Abcam; 1:200). Hoechst^+^NeuN^+^ and Hoechst^+^Olig2^+^ populations were directly sorted into RLT Lysis buffer (#74004, Qiagen) and further processed according to the manufacturer’s instructions.

### High Dimensional Analysis

Raw flow cytometry data were pre-processed in FlowJo (Tree Star Inc.). The high dimensional analysis was performed in Rstudio version 4.0.1 according to the analysis pipeline described previously^91^. In brief, UMAPs ^20^ were generated using the package *umap* version 0.2.7.0, and FlowSOM clustering ^21^ was overlaid on the dimensionality reduction maps. Frequency plots were generated using the *ggplot2* package version 3.3.5, and heatmaps were generated using the *pheatmap* package version 1.0.12.

### Statistical Analysis

Statistical analysis was carried out using GraphPad Prism 8 (GraphPad Software, Inc.). Statistical significance of *in vivo* experiments (clinical scores over time) was determined with a two-way analysis of variance (ANOVA) with Bonferroni’s post-test, Tukey’s or Sidak’s multiple comparisons. Comparisons between two groups were performed with a two-tailed, unpaired *t* test or Mann-Whitney test or Kruskal–Wallis test, corrected Dunn’s test. Statistical significance of western blot quantitative data was evaluated by one-way ANOVA with Bonferroni’s multiple comparison post-hoc test. Statistical analysis of CNS infiltration of different cell types between different groups was carried out in Rstudio (version 4.0.1) using *t* tests. P values below 0.05 were considered as significant and are indicated by asterisks (*P < 0.05, **P < 0.01, ***P < 0.001; ****P < 0.0001) or numerical values on the respective graphs. *N* represents the number of independent biological replicates. In all cases, data are shown as mean (±SEM), unless otherwise indicated. No statistical methods were used to predetermine sample size. Sample sizes were chosen according to previous studies in the field^75^.

## Data Availability

All sequencing data will be publicly available upon publication.

## Acknowledgements

We thank Lucy Robinson from Insight Editing London for manuscript editing; Aakriti Sethi, Sabrina Marti, Mirjam Lutz, Nicole Puertas and André Fonseca da Silva for technical support; Hubert Rehrauer, Lennart Opitz, Andreia Cabral de Gouvea, Doris Popovic, and Timothy Sykes for technical and computational advice on sequencing experiments; the Center for Microscopy and Image Analysis, University of Zurich (ZMB) and the Functional Genomics Center Zurich (FGCZ) for providing equipment. This work has received funding from the Swiss National Science Foundation (Ambizione grant PZ00P3_193330 to S.M. and 733 310030_170320, 310030_188450, and CRSII5_183478 to B.B.), European Research Council (ERC) under the European Union’s Horizon 2020 research and innovation program (grant agreement No 882424 to B.B.), Research Talent Development Fund zur Förderung des akademischen Nachwuchses (FAN) (to S.M.), Forschungskredit Postdoc fellowship University of Zurich grant number: K-41302-11-01 (to S.M.), Dr. Wilhelm Hurka Foundation (to B.B and S.M.) and Swiss Multiple Sclerosis Society (to S.M. and F.I.). F.I. received a PhD fellowship from Studienstiftung des deutschen Volkes.

## Contributions

M.A. designed and performed all mouse experiments including data analysis. S.T. and D.D.F. assisted with single-nucleus RNA sequencing experiments. S.S., M.G. and T.L.M.C. generated primary mouse cultures. P.E. generated multiplexed fluorescence *in situ* hybridization data. D.D.F, S.M., S.T., F.R. helped with mouse experiments. D.K. provided human samples. F.I., E.F., S.M., D.D.F. performed bioinformatics analyses. B.S. and C.W. performed tissue immunostainings. S.M., B.B, D.D.F., C.W., M.G., S.S., S.T. and F.L.H. provided scientific and intellectual input. All authors approved the manuscript. M.A., S.M. and B.B. wrote the manuscript. S.M. and B.B. conceptualized, jointly supervised and funded the study.

## Supplementary data

**Supplementary Table 1**: Differential gene expression between *Nestin^Cre/+^Il12rb2^fl/fl^* and *Il12rb2*^fl/fl^ mice in the steady-state.

**Supplementary Table 2:** Differential gene expression between *Nestin^Cre/+^Il12rb2^fl/fl^* and *Il12rb2*^fl/fl^ mice in EAE.

**Supplementary Table 3**: Pathway analysis (KEGG) of DEGs between *Nestin^Cre/+^Il12rb2^fl/fl^* versus *Il12rb2^fl/fl^* mice in EAE.

**Supplementary Table 4:** Receptor-ligand interactions differentially coexpressed between *Nestin^Cre/+^Il12rb2^fl/fl^* and *Il12rb2^fl/fl^* mice in EAE.

**Supplementary Table 5:** Differentially expressed genes in IL-12 (100 ng/μl) treated primary murine neuronal cultures compared to unstimulated.

## Extended data figures and figure legends

**Extended Data Fig.1:**
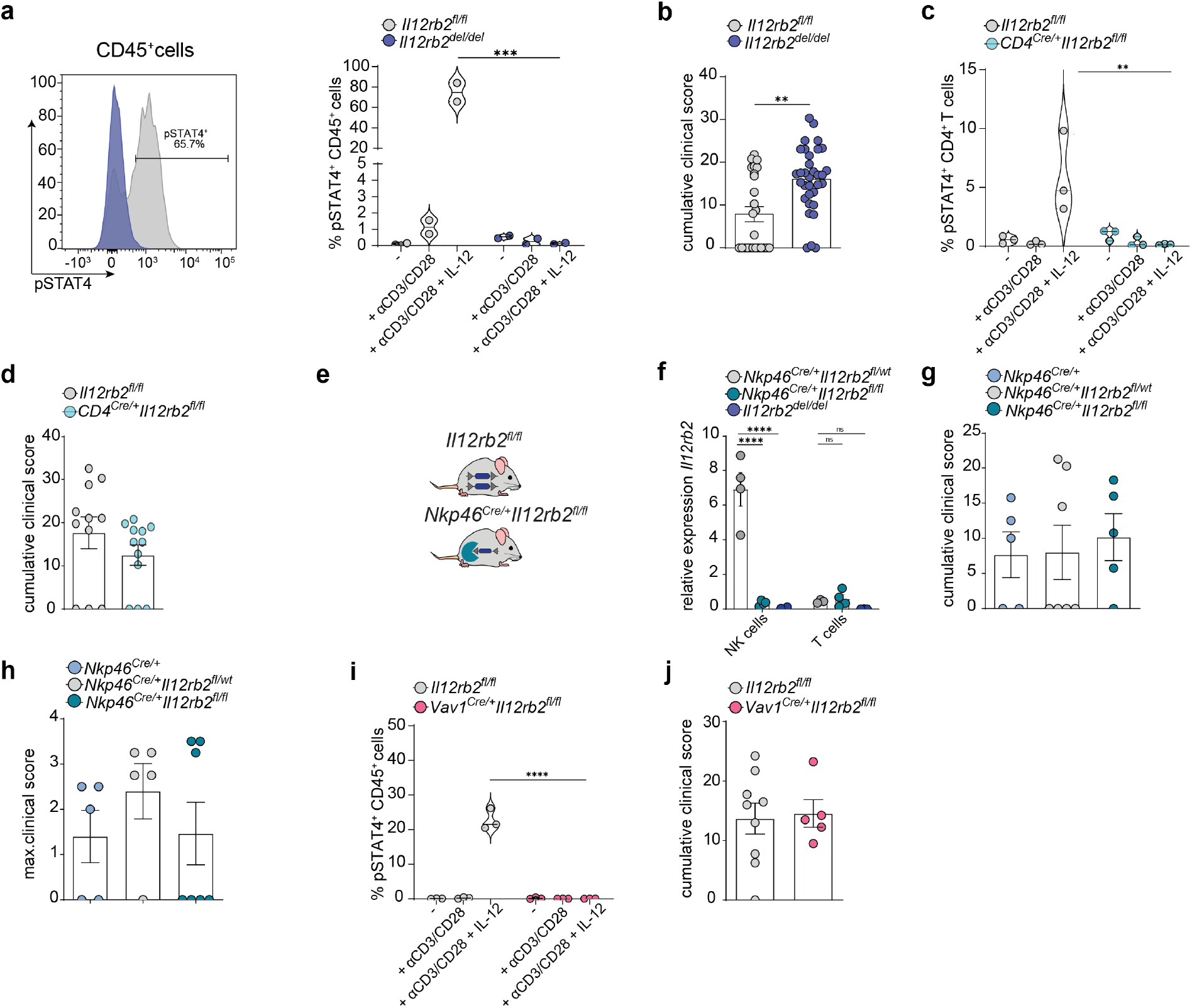
Loss of IL-12 signalling across the leukocyte compartment does not influence susceptibility to EAE. (a) Representative histogram and frequencies of pSTAT4^+^ CD45^+^ cells upon IL-12 stimulation of splenocytes isolated from *Il12rb2^del/del^* and *Il12rb2^fl/fl^* mice. Data represent two experiments (n=2 per group). (b) Cumulative disease scores of MOG-CFA induced EAE in *Il12rb2^del/del^* (n=33) and *Il12rb2^fl/fl^* mice (n=25). Data pooled from four independent experiments using both male and female mice. (c) Frequency of pSTAT4^+^CD4^+^cells upon IL-12 stimulation of splenocytes isolated from steady-state *CD4^Cre/+^Il12rb2^fl/fl^* and littermate controls. Data represent one experiment (n=3 per group). (d) Cumulative clinical score of MOG-CFA induced EAE in *CD4^Cre/+^Il12rb2^fl/fl^* (n=12) and littermate controls (n=11). Data are pooled from two independent experiments. (e) *Nkp46^Cre/+^* mice were crossed to *Il12rb2^fl/fl^* mice to generate *Nkp46^Cre/+^Il12rb2^fl/fl^* mice, which lack *Il12rb2* in NK cells. (f) *Il12rb2* mRNA expression of NK cells and CD3^+^ T cells isolated by FACS from the spleens of *Nkp46^Cre/+^Il12rb2^fl/fl^* (n=4), littermate *Nkp46^Cre/+^Il12rb2^fl/wt^* controls (n=4) and *Il12rb2^del/del^* (n=3) mice at 17 dpi. Data represent one experiment. (g) Cumulative and (h) maximal clinical scores of MOG-CFA induced EAE in *Nkp46^Cre/+^* (n=5), *Nkp46^Cre/+^Il12rb2^fl/wt^* (n=5) and *Nkp46^Cre/+^Il12rb2^fl/fl^* mice (n=7). Data represent one experiment. (i) Frequency of pSTAT4^+^CD45^+^cells upon IL-12 stimulation of splenocytes isolated from steady-state *Vav1^Cre/+^Il12rb2^fl/fl^* and littermate controls. Data represent one experiment (n=3 per group). (j) Cumulative disease scores of MOG-CFA induced EAE in *Vav1^Cre/+^Il12rb2^fl/fl^* (n=5) and littermate controls (n=9). The data are representative of two independent experiments. Each symbol represents one mouse. Data are shown as mean ± SEM.In (a), (c), (f) and (i) statistical significance was evaluated by two-way ANOVA with Bonferroni’s post hoc test; an unpaired Student’s t test was used to compare the means in (b), (d) and (g); a non-parametric Mann-Whitney test in (h). * P < 0.05, ** P < 0.01,*** P < 0.001, ****P < 0.0001, ns=not significant.

**Extended Data Fig.2:**
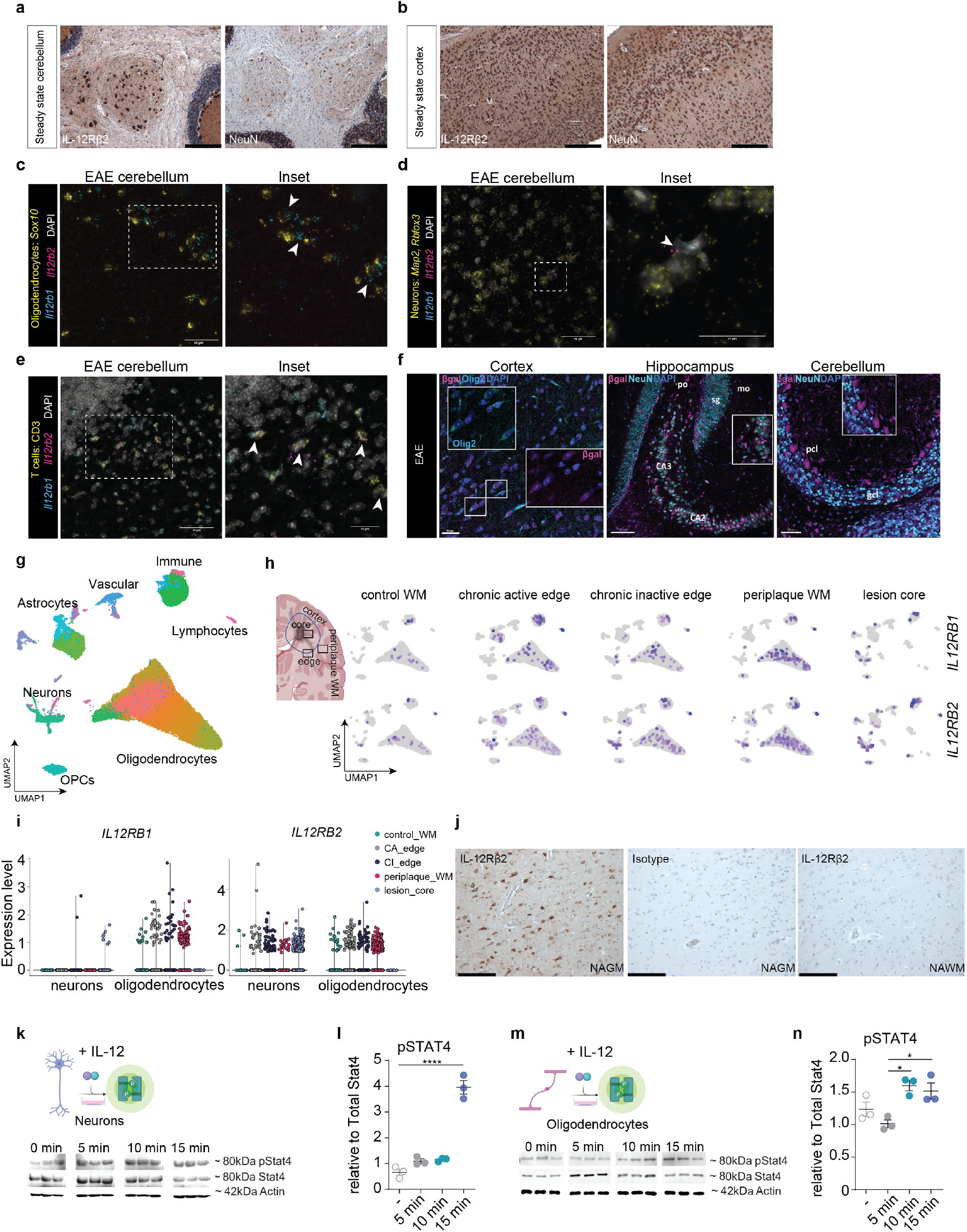
Murine and human neuroectodermal cells harbour the molecular repertoire to execute IL-12 signalling in steady-state and disease. (a-b) Representative immunohistochemistry for IL-12Rβ2^+^and NeuN^+^ cells in the cerebellum and cortex of the steady-state C57BL/6 wt mouse brain; scale bar= 200 μm. (c-e) Representative images of multiplexed single-molecule RNA fluorescence in situ hybridization (RNAscope^®^) staining of EAE C57BL/6 wt mouse brain tissue using target probes for oligodendrocytes (*Sox10*), neurons (*Map2, Rbfox3*) and T cells (CD3). Cell markers were co-stained with probes for *Il12rb1* and *Il12rb2*, as highlighted by arrowheads. Scale bars = 25 μm (insets) or 50 μm. (f) Immunostaining of β-gal^+^Olig2^+^ cells (cortex) and β-gal^+^NeuN^+^ cells (hippocampus and cerebellum) in EAE *Il12rb2^LacZ^* reporter mice (n≥4mice, ≥5 sections per mouse). Scale bars = 80 μm or 100 μm. (g) Single-nucleus RNAseq of n=66,432 nuclei from white matter areas of five patients with progressive MS and three age- and sex-matched non affected, non-dementia controls, analyzed by Absinta *et* al., 2021^18^ (accessed at GSE180759). UMAP plot displaying all identified cell clusters, sorted by cell type (OPCs: oligodendrocyte progenitor cells). (h) Schematic illustration of MS lesion, created with Biorender.com (left). UMAP plots displaying *IL12RB1*- and *IL12RB2*-expressing cells, sorted by pathological condition (right). (i) Violin plots displaying *IL12RB1* and *IL12RB2* transcripts in neurons and oligodendrocytes, split by pathological condition. (j) Brain sections from patients with MS were stained with antibodies against human IL-12Rβ2 or total rabbit IgG for immunohistochemistry; scale bar= 200 μm [normal appearing grey matter (NAGM), normal appearing white matter (NAWM)]. (k) Whole cell protein lysates were prepared from dissociated cerebellar neurons at 14 days *in vitro* following 5, 10 and 15 min stimulation with IL-12 (100 ng/mL); western blot analysis was performed for total Stat4 normalized to total actin level, and phospho-Stat4 (Tyr693) normalized to total Stat4 levels. (l) Fold change in pSTAT4 (relative to total Stat4) protein expression upon IL-12 engagement in neurons. (m) Whole cell protein lysates were prepared from primary oligodendrocyte cultures at 4 days in vitro following 5, 10 and 15 min stimulation with IL-12 (100 ng/mL); western blot analysis was performed for levels of total Stat4 normalized to total actin level, and pStat4 (Tyr693) normalized to total Stat4 levels. (n) Quantification of IL-12-driven induction of pSTAT4 (relative to total Stat4) in stimulated oligodendrocytes. In (k-l) and (n-o) data are shown as mean ± SD. Statistical significance was evaluated by one-way ANOVA with Bonferroni’s multiple comparison post-hoc test; * P ≤ 0.05, ** P < 0.01, *** P < 0.001, P < 0.0001****. GCL, granule cell layer; PCL, Purkinje cell layer; sg (DG-sg), dentate gyrus, granule cell layer; po (DG-po), dentate gyrus, polymorph layer; mo (DG-mo), dentate gyrus, molecular layer.

**Extended Data Fig.3:**
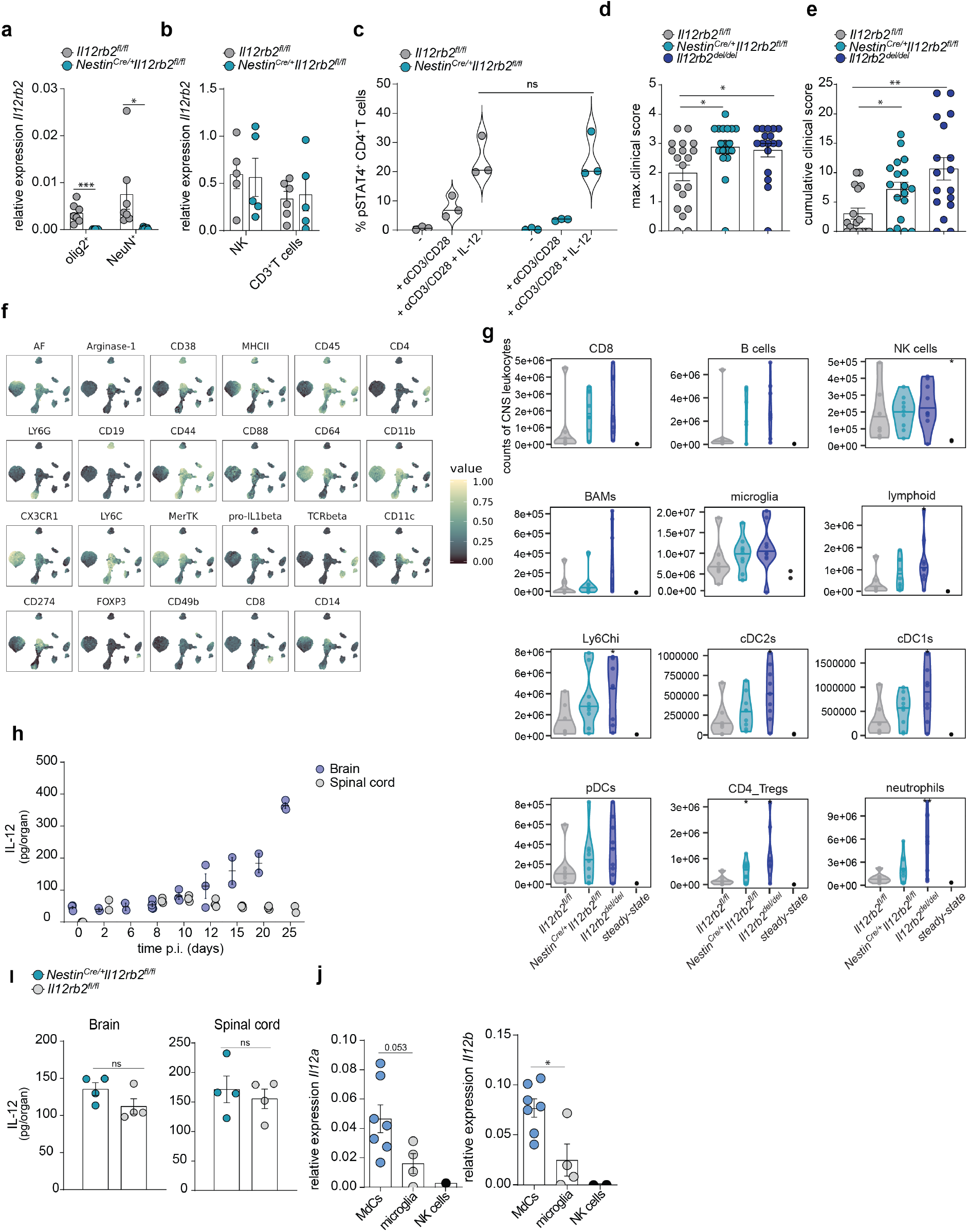
IL-12 engagement in neurons and oligodendrocytes protects against neuroinflammation. (a) *Il12rb2* expression in oligodendrocyte nuclei (Hoechst^+^Olig2^+^) and neuronal nuclei (Hoechst^+^NeuN^+^) isolated by FACS from the CNS of *Nestin^Cre/+^Il12rb2f^l/fl^* (n=7) and control mice (n=7) at peak EAE (13 dpi). Data represent two experiments shown as mean ± SEM. Unpaired two-tailed Mann-Whitney test. * P < 0.05, ** P < 0.01, *** P < 0.001. (b) *Il12rb2* mRNA expression in NK cells and CD3^+^ T cells isolated by FACS from the CNS of *Nestin^Cre/+^Il12rb2^fl/fl^* and littermate controls at late stage EAE (17 dpi) (n=3 to 6 per group). Pooled data from two independent experiments shown as mean ± SEM. (c) Frequency of pSTAT4^+^ CD4^+^cells upon IL-12 stimulation of splenocytes from steady-state *Nestin^Cre/+^Il12rb2^fl/fl^* (n=3) mice and littermate controls (n=3). Data represent one experiment. Shown as mean ± SEM. Two-way ANOVA with Bonferroni’s post test. NS= not significant. (d) Individual maximum disease scores and (e) Cumulative clinical score of MOG-CFA induced EAE in *Nestin^Cre/+^Il12rb2^fl/fl^* (n=18), littermate *Il12rb2^fl/fl^* control (n=19) and *Il12rb2^del/del^* (n=17) mice. Data pooled from four independent experiments. Each symbol represents one mouse. Data represent mean ± SEM. Unpaired Student’s t test. *P < 0.05, **P < 0.01. (f) Spectral cytometry panel markers overlaid onto UMAP from Fig. 3c of CNS leukocytes from *Nestin^Cre/+^Il12rb2^fl/fl^* (n=11), *Il12rb2^fl/fl^* (n=7), *Il12rb2^del/del^* (n=9) and steady-state mice at peak EAE (13 dpi). (g) Cell counts of leukocytes from (f); *t*-test. (h) Kinetics of IL-12 (IL-12/23p40) concentration in the brain and spinal cord during MOG-CFA induced EAE in C57BL/6 mice. Each data point represents an individual biological replicate (n=2 or n=3 per time point). (i) *Nestin^Cre/+^Il12rb2^fl/fl^* (n=4) and littermate controls (n=4) were sacrificed at 10 dpi and the levels of IL-12 were measured in lysates of the brain and spinal cord. Each data point represents an individual biological replicate. Data represent mean ± SEM. An unpaired Student’s t test was used to compare the means. NS= not significant. (j) *Il12a* and *Il12b* mRNA expression in MdCs (CD45^+^CD11b^+^Ly6G^-^ Ly6C^+^CD44^+^CX3CR1^-^), microglia (CD45^low^CD11b^+^CX3CR1^+^), and NK cells (negative control) isolated by FACS from C57BL/6 wt mouse brain tissue at disease onset (10 dpi). Each data point represents a biological replicate. Shown as mean ± SEM. Unpaired *t* test. NS= not significant.

**Extended Data Fig.4:**
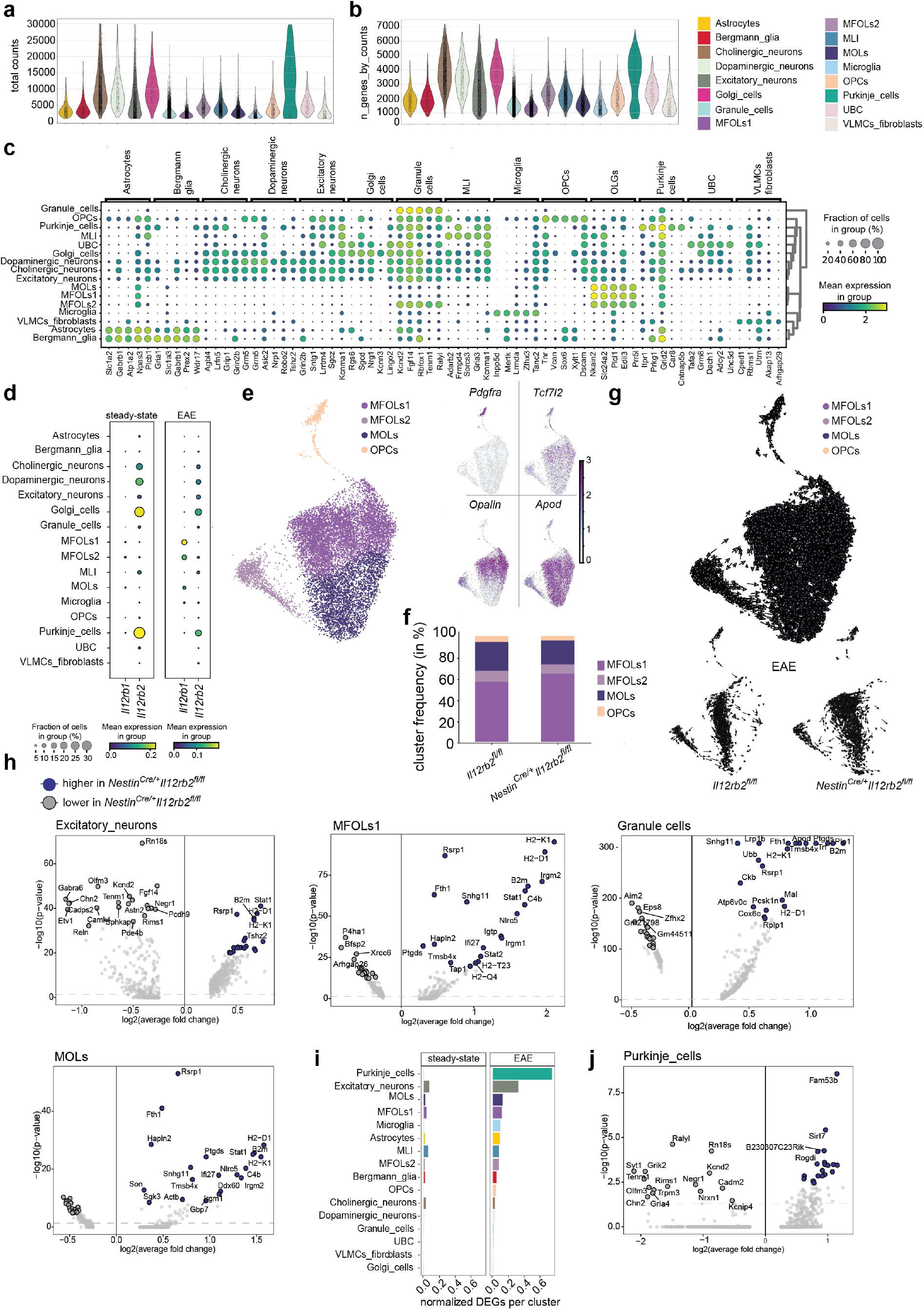
Direct IL-12 engagement of neuronal cells mediates neuroprotection. (a-b) Nuclei were isolated from the CNS of *Il12rb2^fl/fl^* and *Nestin^Cre/+^Il12rb2^fl/fl^* male mice (n=1 per genotype, 8-12 weeks old) at the steady-state and early onset of EAE (10 dpi, clinical score<1). The Hoechst^+^ fraction was FANS-selected and used for single-nucleus RNA sequencing. Violin plots depicting the distribution of (a) number of total counts and (b) number of genes by counts detected in each cluster colour coded as in Fig. 4b. (c) Dot plot depicting the top 5 differentially expressed genes (DEGs) for each cluster and associated cluster labelling. Dot size corresponds to the percentage of nuclei expressing marker genes across all clusters and the color shows the average expression level. (d) Dot plots depicting the expression of *Il12rb1* and *Il12rb2* for each cluster and associated cluster labelling. Dot size corresponds to the percentage of nuclei expressing the gene in each cluster, and the colour represents the average expression level. (e) Subclustering (UMAP plot) and annotation for the oligodendrocyte lineage (OL) based on DEGs in each cluster showing the continuous developmental process from precursor to mature cells states with feature plots displaying marker genes of oligodendrocyte cell maturation states, as described in Marques *et* al, 2016. (f) Distribution of OL lineage cellular proportions across genotype in EAE. (g) OL lineage UMAP [as defined in (e)] showing the differentiation trajectory from OPCs to mature oligodendrocytes. Embedded arrows show RNA velocity derived from the dynamic model (top part). Velocities derived from the dynamic model split by genotype in EAE (bottom part). (h) Volcano plots displaying genes that are up-(blue) or downregulated (grey) in the inflamed *Nestin^Cre,/+^Il12rb2^fl,/fl^* CNS for the highlighted neuronal or oligodendrocyte clusters. Dashed lines indicate significance thresholds (adjusted p-value < 0.05) used when identifying DEGs. (i) Number of DEGs, normalized per cluster size between *Nestin^Cre,/+^Il12rb2^fl/fl^* and *Il12rb2^f/fl^* mice across the major cell types in steady-state and during EAE. (j) Volcano plot displaying genes that are up- (blue) or downregulated (grey) in the inflamed *Nestin^Cre/+^Il12rb2^fl/fl^* CNS for Purkinje cells. Dashed lines indicate significance thresholds (adjusted p-value < 0.05) used when identifying DEGs.

**Extended Data Fig.5:**
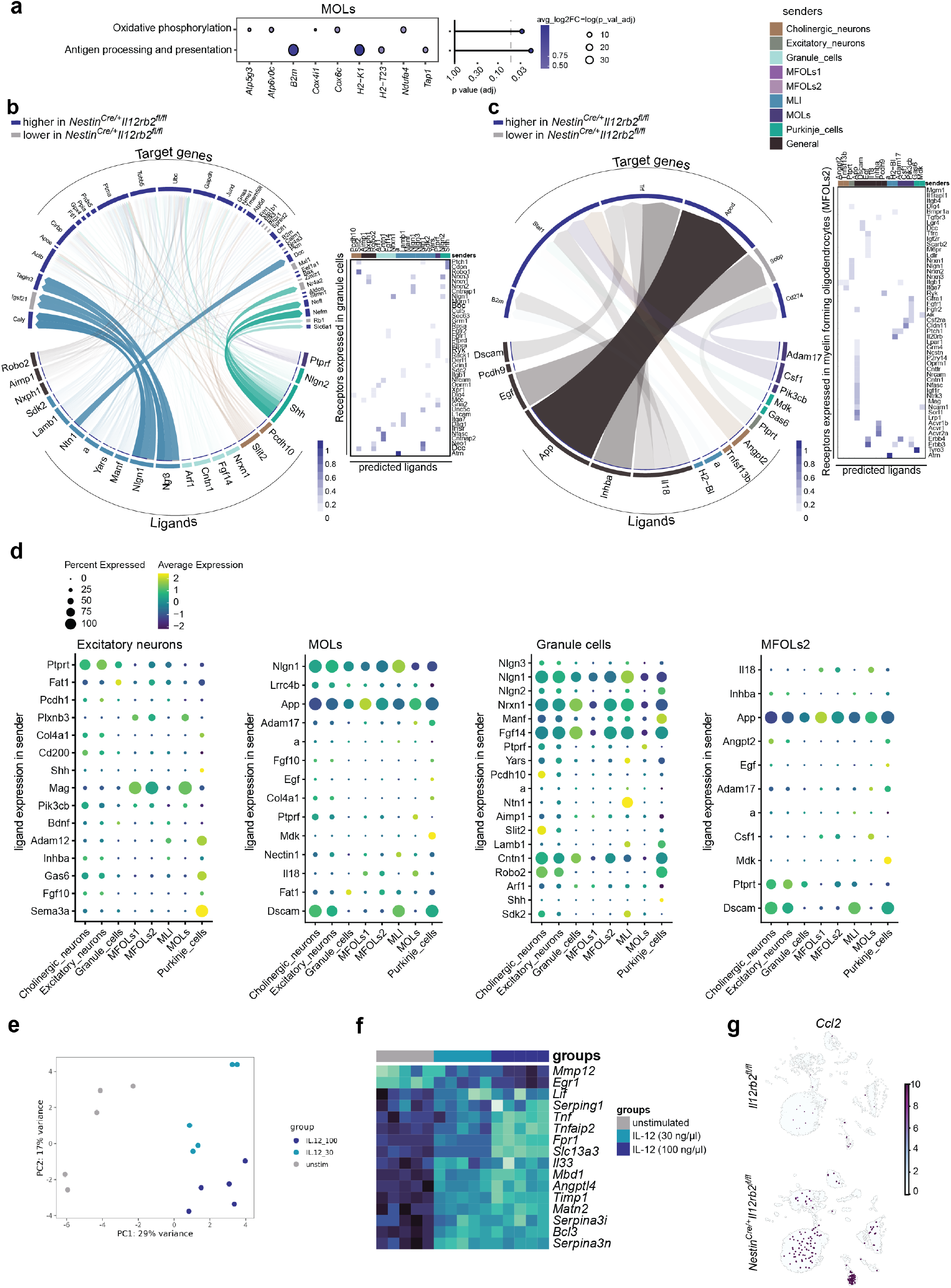
IL-12 induces neuroprotection and sustains trophic factor release in the inflamed CNS. (a) Pathway analysis (KEGG) of DEGs from MOLs of EAE *Nestin^Cre/+^ Il12rb2^fl/fl^* versus *Il12rb2^fl/fl^* mice. Top2 KEGG pathways with g:SCS-corrected p-values < 0.05 are shown. (b) Receptor-ligand interaction analysis (NicheNet). Circos plot (left) showing links between unique ligands from cholinergic neurons, excitatory neurons, Purkinje cells, MLIs, MOLs, MFOLs1 and MFOLs2 (senders; ribbon colour indicates subcluster of origin for each ligand) or general ligands (multiple sender cells; ribbon colour black) and predicted associated DE genes in granule cells (*Nestin^Cre/+^Il12rb2^fl/fl^* versus *Il12rb2^fl/fl^* EAE, blue: higher in *Nestin^Cre/+^Il12rb2^fl/fl^*, grey: lower in *Nestin*^*Cre*/+^*Il12rb2^fl/fl^*) potentially targeted by the ligand-receptor pairs. Transparency indicates interaction strength. Heatmap (right) displaying potential receptors expressed in granule cells associated with each predicted ligand. (c) Circos plot (left) showing links between unique ligands from cholinergic neurons, excitatory neurons, granule cells, Purkinje cells, MLIs, MFOLs1 and MOLs (senders; ribbon colour indicates subcluster of origin for each ligand) or general ligands (multiple sender cells; ribbon colour black) and predicted associated DE genes in MFOLs2 (*Nestin^Cre/+^Il12rb2^fl/fl^* versus *Il12rb2^fl/fl^* EAE, blue: higher in *Nestin^Cre/+^Il12rb2^fl/fl^*, grey: lower in *Nestin^Cre/+^Il12rb2^fl/fl^*) potentially targeted by the ligand-receptor pairs. Heatmap (right) displaying potential receptors expressed in MFOLs2 associated with each predicted ligand. In (b) and (c) the ribbon thickness is proportional to the ligand’s regulatory potential. (d) Dot plots depicting the expression of the predicted ligands (in sender) for the following recipient clusters: excitatory neurons, MOLs, granule cells and MFOLs2 (left to right). Dot size corresponds to the percentage of nuclei expressing the gene in each cluster, and the color represents the average expression level. (e) Primary neuronal cultures were activated with IL-12 (30 ng/μl and 100 ng/μl) for 18 hours and harvested for RNAseq (n=5 technical replicates per group). Principal Component Analysis (PCA) of bulk RNAseq shows a dose-dependent effect of IL-12 on neurons. (f) Heatmap depicting significantly DEGs (FDR< 0.05) between untreated and IL-12-activated neurons (30 ng/μl and 100 ng/μl). (g) Feature plots displaying the expression of *Ccl2* across clusters in EAE.

## References

1. Reich, D. S., Lucchinetti, C. F. & Calabresi, P. A. Multiple Sclerosis. https://doi.org/10.1056/NEJMra1401483 378, 169–180 (2018).

2. Kuhlmann, T. et al. An updated histological classification system for multiple sclerosis lesions. Acta Neuropathol. 133, 13–24 (2017).

3. Trinchieri, G. Interleukin-12: A Proinflammatory Cytokine with Immunoregulatory Functions that Bridge Innate Resistance and Antigen-Specific Adaptive Immunity. Annu. Rev. Immunol. 13, 251–276 (1995).

4. Oppmann, B. et al. Novel p19 Protein Engages IL-12p40 to Form a Cytokine, IL-23, with Biological Activities Similar as Well as Distinct from IL-12. Immunity 13, 715–725 (2000).

5. Cua, D. J. et al. Interleukin-23 rather than interleukin-12 is the critical cytokine for autoimmune inflammation of the brain. Nature (2003) doi:10.1038/nature01355.

6. Langrish, C. L. et al. IL-23 drives a pathogenic T cell population that induces autoimmune inflammation. 201, 233–240 (2005).

7. Becher, B., Durell, B. G. & Noelle, R. J. Experimental autoimmune encephalitis and inflammation in the absence of interleukin-12. J. Clin. Invest. (2002) doi:10.1172/JCI200215751.

8. Gran, B. et al. IL-12p35-Deficient Mice Are Susceptible to Experimental Autoimmune Encephalomyelitis: Evidence for Redundancy in the IL-12 System in the Induction of Central Nervous System Autoimmune Demyelination. J. Immunol. 169, 7104–7110 (2002).

9. Segal, B. M. et al. Repeated subcutaneous injections of IL12/23 p40 neutralising antibody, ustekinumab, in patients with relapsing-remitting multiple sclerosis: a phase II, double-blind, placebo-controlled, randomised, dose-ranging study. Lancet Neurol. 7, 796–804 (2008).

10. Grossman, I. et al. Pharmacogenetics of glatiramer acetate therapy for multiple sclerosis reveals drug-response markers. Pharmacogenet. Genomics 17, 657–666 (2007).

11. Milosevic, E. et al. Higher expression of IL-12Rβ2 is associated with lower risk of relapse in relapsing-remitting multiple sclerosis patients on interferon-β1b therapy during 3-year follow-up. J. Neuroimmunol. 287, 64–70 (2015).

12. Zwicky, P. et al. IL-12 regulates type 3 immunity through interfollicular keratinocytes in psoriasiform inflammation. Sci. Immunol. 6, 9012 (2021).

13. Goverman, J. Autoimmune T cell responses in the central nervous system. Immunology 9, 393–407 (2009).

14. Utz, S. G. et al. Early Fate Defines Microglia and Non-parenchymal Brain Macrophage Development. Cell 181, 557–573.e18 (2020).

15. Kozareva, V. et al. A transcriptomic atlas of mouse cerebellar cortex comprehensively defines cell types. Nat. 2021 5987879 598, 214–219 (2021).

16. Habib, N. et al. Div-Seq: Single-nucleus RNA-Seq reveals dynamics of rare adult newborn neurons. Science (80-.). 353, 925–928 (2016).

17. Schneeberger, S. et al. The neuroinflammatory interleukin-12 signaling pathway drives Alzheimer’s disease-like pathology by perturbing oligodendrocyte survival and neuronal homeostasis. bioRxiv 2021.04.25.441313 (2021) doi:10.1101/2021.04.25.441313.

18. Absinta, M. et al. A lymphocyte–microglia–astrocyte axis in chronic active multiple sclerosis. Nat. 2021 1–6 (2021) doi:10.1038/s41586-021-03892-7.

19. Tronche, F. et al. Disruption of the glucocorticoid receptor gene in the nervous system results in reduced anxiety. Nat. Genet. 23, 99–103 (1999).

20. McInnes, L., Healy, J. & Melville, J. UMAP: Uniform Manifold Approximation and Projection for Dimension Reduction. arXiv Prepr. (2018).

21. Van Gassen, S. et al. FlowSOM: Using self-organizing maps for visualization and interpretation of cytometry data. Cytom. Part A 87, 636–645 (2015).

22. Komuczki, J. et al. Fate-Mapping of GM-CSF Expression Identifies a Discrete Subset of Inflammation-Driving T Helper Cells Regulated by Cytokines IL-23 and IL-1β. Immunity 50, 1289–1304.e6 (2019).

23. Mrdjen, D. et al. High-Dimensional Single-Cell Mapping of Central Nervous System Immune Cells Reveals Distinct Myeloid Subsets in Health, Aging, and Disease. Immunity 48, 380–395.e6 (2018).

24. Habib, N. et al. Massively parallel single-nucleus RNA-seq with DroNc-seq. Nat. Methods 14, 955–958 (2017).

25. Marques, S. et al. Oligodendrocyte heterogeneity in the mouse juvenile and adult central nervous system. Science (80-.). 352, 1326–1329 (2016).

26. Zeisel, A. et al. Molecular Architecture of the Mouse Nervous System. Cell 174, 999–1014.e22 (2018).

27. Saunders, A. et al. Molecular Diversity and Specializations among the Cells of the Adult Mouse Brain. Cell 174, 1015–1030.e16 (2018).

28. Keller, D., Erö, C. & Markram, H. Cell densities in the mouse brain: A systematic review. Frontiers in Neuroanatomy (2018) doi:10.3389/fnana.2018.00083.

29. Erö, C., Gewaltig, M. O., Keller, D. & Markram, H. A cell atlas for the mouse brain. Front. Neuroinform. (2018) doi:10.3389/fninf.2018.00084.

30. Franklin, R. J. M. & Ffrench-Constant, C. Regenerating CNS myelin — from mechanisms to experimental medicines. Nat. Rev. Neurosci. 2017 18 12 18, 753–769 (2017).

31. Marques, S. et al. Oligodendrocyte heterogeneity in the mouse juvenile and adult central nervous system. Science (80-.). 352, 1326–1329 (2016).

32. Golan, N. et al. Genetic Deletion of Cadm4 Results in Myelin Abnormalities Resembling Charcot-Marie-Tooth Neuropathy. J. Neurosci. 33, 10950–10961 (2013).

33. Lammert, C. R. et al. AIM2 inflammasome surveillance of DNA damage shapes neurodevelopment. Nat. 2020 5807805 580, 647–652 (2020).

34. Kenigsbuch, M. et al. A shared disease-associated oligodendrocyte signature among multiple CNS pathologies. Nat. Neurosci. 2022 1–11 (2022) doi:10.1038/s41593-022-01104-7.

35. Browaeys, R., Saelens, W. & Saeys, Y. NicheNet: modeling intercellular communication by linking ligands to target genes. Nat. Methods 2019 17 2 17, 159–162 (2019).

36. Chechneva, O. V. et al. A Smoothened receptor agonist is neuroprotective and promotes regeneration after ischemic brain injury. Cell Death Dis. 2014 510 5, e1481–e1481 (2014).

37. Giovane, A. Del & Ragnini-Wilson, A. Molecular Sciences Targeting Smoothened as a New Frontier in the Functional Recovery of Central Nervous System Demyelinating Pathologies. (2018) doi:10.3390/ijms19113677.

38. Cuttler, K., Hassan, M., Carr, J., Cloete, R. & Bardien, S. Emerging evidence implicating a role for neurexins in neurodegenerative and neuropsychiatric disorders. Open Biol. 11, (2021).

39. Kattimani, Y. & Veerappa, A. M. Dysregulation of NRXN1 by mutant MIR8485 leads to calcium overload in pre-synapses inducing neurodegeneration in Multiple sclerosis. Mult. Scler. Relat. Disord. 22, 153–156 (2018).

40. Scalabrino, G. Epidermal Growth Factor in the CNS: A Beguiling Journey from Integrated Cell Biology to Multiple Sclerosis. An Extensive Translational Overview. Cell. Mol. Neurobiol. 2020 424 42, 891–916 (2020).

41. Scalabrino, G. New Epidermal-Growth-Factor-Related Insights Into the Pathogenesis of Multiple Sclerosis: Is It Also Epistemology? Front. Neurol. 12, 2215 (2021).

42. Fredrickx, E. et al. Ablation of neuronal ADAM17 impairs oligodendrocyte differentiation and myelination. Glia 68, 1148–1164 (2020).

43. Nicoletti, F. et al. Prevention of clinical and histological signs of MOG-induced experimental allergic encephalomyelitis by prolonged treatment with recombinant human EGF. J. Neuroimmunol. 332, 224–232 (2019).

44. Brinkmann, B. G. et al. Neuregulin-1/ErbB Signaling Serves Distinct Functions in Myelination of the Peripheral and Central Nervous System. Neuron (2008) doi:10.1016/j.neuron.2008.06.028.

45. Akkermann, R. et al. The TAM receptor Tyro3 regulates myelination in the central nervous system. Glia 65, 581–591 (2017).

46. Jha, S. et al. The Inflammasome Sensor, NLRP3, Regulates CNS Inflammation and Demyelination via Caspase-1 and Interleukin-18. J. Neurosci. 30, 15811–15820 (2010).

47. Haile, Y. et al. Granzyme B-inhibitor serpina3n induces neuroprotection in vitro and in vivo. (2012) doi:10.1186/s12974-015-0376-7.

48. Burtscher, J., Mallet, R. T., Burtscher, M. & Millet, G. P. Hypoxia and brain aging: Neurodegeneration or neuroprotection? Ageing Res. Rev. 68, 101343 (2021).

49. Leibinger, M., Andreadaki, A. & Fischer, D. Role of mTOR in neuroprotection and axon regeneration after inflammatory stimulation. Neurobiol. Dis. 46, 314–324 (2012).

50. Chan, S. H. et al. Induction of interferon gamma production by natural killer cell stimulatory factor: characterization of the responder cells and synergy with other inducers. J. Exp. Med. 173, 869–879 (1991).

51. Gazzinelli, R. T., Hieny, S., Wynn, T. A., Wolf, S. & Sher, A. Interleukin 12 is required for the T-lymphocyte-independent induction of interferon gamma by an intracellular parasite and induces resistance in T-cell-deficient hosts. Proc. Natl. Acad. Sci. 90, 6115–6119 (1993).

52. Tuzlak, S. et al. Repositioning TH cell polarization from single cytokines to complex help. Nat. Immunol. 2021 2210 22, 1210–1217 (2021).

53. Teng, M. W. L. et al. IL-12 and IL-23 cytokines: From discovery to targeted therapies for immune-mediated inflammatory diseases. Nature Medicine (2015) doi:10.1038/nm.3895.

54. Tait Wojno, E. D., Hunter, C. A. & Stumhofer, J. S. The Immunobiology of the Interleukin-12 Family: Room for Discovery. Immunity vol. 50 851–870 (2019).

55. Becher, B., Spath, S. & Goverman, J. Cytokine networks in neuroinflammation. Nat. Rev. Immunol. 2016 171 17, 49–59 (2016).

56. Kaufmann, M. et al. Identification of early neurodegenerative pathways in progressive multiple sclerosis. Nat. Neurosci. 2022 1–12 (2022) doi:10.1038/s41593-022-01097-3.

57. Jin, K. et al. Cerebral neurogenesis is induced by intranasal administration of growth factors. Ann. Neurol. 53, 405–409 (2003).

58. Ruffini, F. et al. Fibroblast growth factor-II gene therapy reverts the clinical course and the pathological signs of chronic experimental autoimmune encephalomyelitis in C57BL/6 mice. Gene Ther. 2001 816 8, 1207–1213 (2001).

59. Jin, K. et al. FGF-2 promotes neurogenesis and neuroprotection and prolongs survival in a transgenic mouse model of Huntington’s disease. Proc. Natl. Acad. Sci. U. S. A. 102, 18189–18194 (2005).

60. Furusho, M., Roulois, A. J., Franklin, R. J. M. & Bansal, R. Fibroblast growth factor signaling in oligodendrocyte-lineage cells facilitates recovery of chronically demyelinated lesions but is redundant in acute lesions. Glia 63, 1714–1728 (2015).

61. Timmer, M. et al. Fibroblast growth factor (FGF)-2 and FGF receptor 3 are required for the development of the substantia nigra, and FGF-2 plays a crucial role for the rescue of dopaminergic neurons after 6-hydroxydopamine lesion. J. Neurosci. 27, 459–471 (2007).

62. Alam, A. et al. Cellular infiltration in traumatic brain injury. J. Neuroinflammation 2020 171 17, 1–17 (2020).

63. Falcão, A. M. et al. Disease-specific oligodendrocyte lineage cells arise in multiple sclerosis. Nature Medicine vol. 24 1837–1844 (2018).

64. Jäkel, S. et al. Altered human oligodendrocyte heterogeneity in multiple sclerosis. Nature vol. 566 543–547 (2019).

65. Schirmer, L. et al. Neuronal vulnerability and multilineage diversity in multiple sclerosis. Nature 573, 75–82 (2019).

66. Wheeler, M. A. et al. MAFG-driven astrocytes promote CNS inflammation. Nature 578, 593–599 (2020).

67. Flügel, A. et al. Neuronal FasL Induces Cell Death of Encephalitogenic T Lymphocytes. Brain Pathol. 10, 353–364 (2006).

68. Di Liberto, G. et al. Neurons under T Cell Attack Coordinate Phagocyte-Mediated Synaptic Stripping. Cell 175, 458–471.e19 (2018).

69. Nakazato, Y., Fujita, Y., Nakazato, M. & Yamashita, T. Neurons promote encephalitogenic CD4+ lymphocyte infiltration in experimental autoimmune encephalomyelitis. Sci. Rep. 10, 1–10 (2020).

70. Liao, G. et al. Enhanced expression of matrix metalloproteinase-12 contributes to Npc1 deficiency-induced axonal degeneration. Exp. Neurol. 269, 67 (2015).

71. Schwenk, F., Baron, U. & Rajewsky, K. A cre-transgenic mouse strain for the ubiquitous deletion of loxP-flanked gene segments including deletion in germ cells. Nucleic Acids Res. 23, 5080–5081 (1995).

72. Luo, L. et al. Optimizing Nervous System-Specific Gene Targeting with Cre Driver Lines: Prevalence of Germline Recombination and Influencing Factors. Neuron 106, 37–65.e5 (2020).

73. Boer, J. de et al. Transgenic mice with hematopoietic and lymphoid specific expression of Cre. Eur. J. Immunol. 33, 314–325 (2003).

74. Sawada, S., Scarborough, J. D., Killeen, N. & Littman, D. R. A lineage-specific transcriptional silencer regulates CD4 gene expression during T lymphocyte development. Cell 77, 917–929 (1994).

75. Mundt, S. et al. Conventional DCs sample and present myelin antigens in the healthy CNS and allow parenchymal T cell entry to initiate neuroinflammation. Sci. Immunol. 4, 8380 (2019).

76. Livak, K. J. & Schmittgen, T. D. Analysis of Relative Gene Expression Data Using Real-Time Quantitative PCR and the 2-ΔΔCT Method. Methods 25, 402–408 (2001).

77. Stamou, M., Grodzki, A. C., van Oostrum, M., Wollscheid, B. & Lein, P. J. Fc gamma receptors are expressed in the developing rat brain and activate downstream signaling molecules upon cross-linking with immune complex. J. Neuroinflammation 2018 151 15, 1–23 (2018).

78. Schneider, C. A., Rasband, W. S. & Eliceiri, K. W. NIH Image to ImageJ: 25 years of image analysis. Nat. Methods 2012 97 9, 671–675 (2012).

79. Mahony, D., Karunaratne, S. & Rothnagel, J. A. Improved detection of lacZ reporter gene expression in transgenic epithelia by immunofluorescence microscopy. Exp. Dermatol. 11, 153–158 (2002).

80. Sommer, C., Straehle, C., Kothe, U. & Hamprecht, F. A. Ilastik: Interactive learning and segmentation toolkit. Proc. - Int. Symp. Biomed. Imaging 230–233 (2011) doi:10.1109/ISBI.2011.5872394.

81. Picelli, S. et al. Full-length RNA-seq from single cells using Smart-seq2. Nat. Protoc. 20l3 9l 9, 171–181 (2014).

82. Robinson, M. D., McCarthy, D. J. & Smyth, G. K. edgeR: a Bioconductor package for differential expression analysis of digital gene expression data. Bioinformatics 26, 139 (2010).

83. Love, M. I., Huber, W. & Anders, S. Moderated estimation of fold change and dispersion for RNA-seq data with DESeq2. Genome Biol. 15, 1–21 (2014).

84. Russo, P. S. T. et al. CEMiTool: A Bioconductor package for performing comprehensive modular co-expression analyses. BMC Bioinformatics 19, 1–13 (2018).

85. Liberzon, A. et al. The Molecular Signatures Database Hallmark Gene Set Collection. Cell Syst. 1, 417–425 (2015).

86. Szklarczyk, D. et al. STRING v11: protein–protein association networks with increased coverage, supporting functional discovery in genome-wide experimental datasets. Nucleic Acids Res. 47, D607–D613 (2019).

87. Wolf, F. A., Angerer, P. & Theis, F. J. SCANPY: Large-scale single-cell gene expression data analysis. Genome Biol. 19, 1–5 (2018).

88. Wolock, S. L., Lopez, R. & Klein, A. M. Scrublet: Computational Identification of Cell Doublets in Single-Cell Transcriptomic Data. Cell Syst. (2019) doi:10.1016/j.cels.2018.11.005.

89. Bergen, V., Lange, M., Peidli, S., Wolf, F. A. & Theis, F. J. Generalizing RNA velocity to transient cell states through dynamical modeling. Nat. Biotechnol. 2020 3812 38, 1408–1414 (2020).

90. Butler, A., Hoffman, P., Smibert, P., Papalexi, E. & Satija, R. Integrating single-cell transcriptomic data across different conditions, technologies, and species A n A ly s I s. Nat. Biotechnol. Vol. 36, (2018).

91. Hartmann, F. J. et al. High-dimensional single-cell analysis reveals the immune signature of narcolepsy. J. Exp. Med. (2016) doi:10.1084/jem.20160897.

